# GenomeCompendium: A database for the integrated analysis of repeats, assembly quality and functional content of complete prokaryotic genomes

**DOI:** 10.64898/2026.08.14.744864

**Authors:** Tiberiu Totu, Garance Jaques, Benjamin Heiniger, Tina Segessemann, Michael Schmid, Marc Bourqui, Adrian Wicki, Jürg E. Frey, Christian H. Ahrens

## Abstract

Microorganisms hold great promise for urgent global needs such as increasing sustainable agricultural production while reducing chemical fertilizer and pesticide use or providing novel classes of antimicrobials/therapeutics. Moving from analyzing microbiome composition to applying synthetic communities and studying their functions requires access to isolates and complete genome sequences. By spanning the frequent repeats, long-read sequencing can resolve complex prokaryotic genomes, yet error-prone short-read assemblies dominate.

We here release the GenomeCompendium, a public database and interactive analysis tool for complete prokaryotic genomes (https://genome-compendium.com/). Using NCBI RefSeq (∼47,000) and GenBank (∼13,000) genomes, we integrated available metadata, GTDB taxonomy and computed features including repeat classes and gene content screening, intragenomic 16S rRNA sequence identity, and biosynthetic gene cluster co-occurrences. Evaluating repeat content and assembly complexity metrics, we identify taxonomic ranks dominated by difficult-to-assemble genomes and show that complex, repeat-rich genomes are more common than previously estimated. By mining metadata, our quality control flags 6.3% of RefSeq assemblies as potentially erroneous or incomplete. As valuable reference for data mining and to track taxonomic coverage, the GenomeCompendium links ∼90 features across genomes, offers downloadable reports and -as unique features-pre-computed proteogenomics databases to improve genome annotations of RefSeq strains and the ability to analyze any uploaded prokaryotic genome.

**Graphical Abstract:** 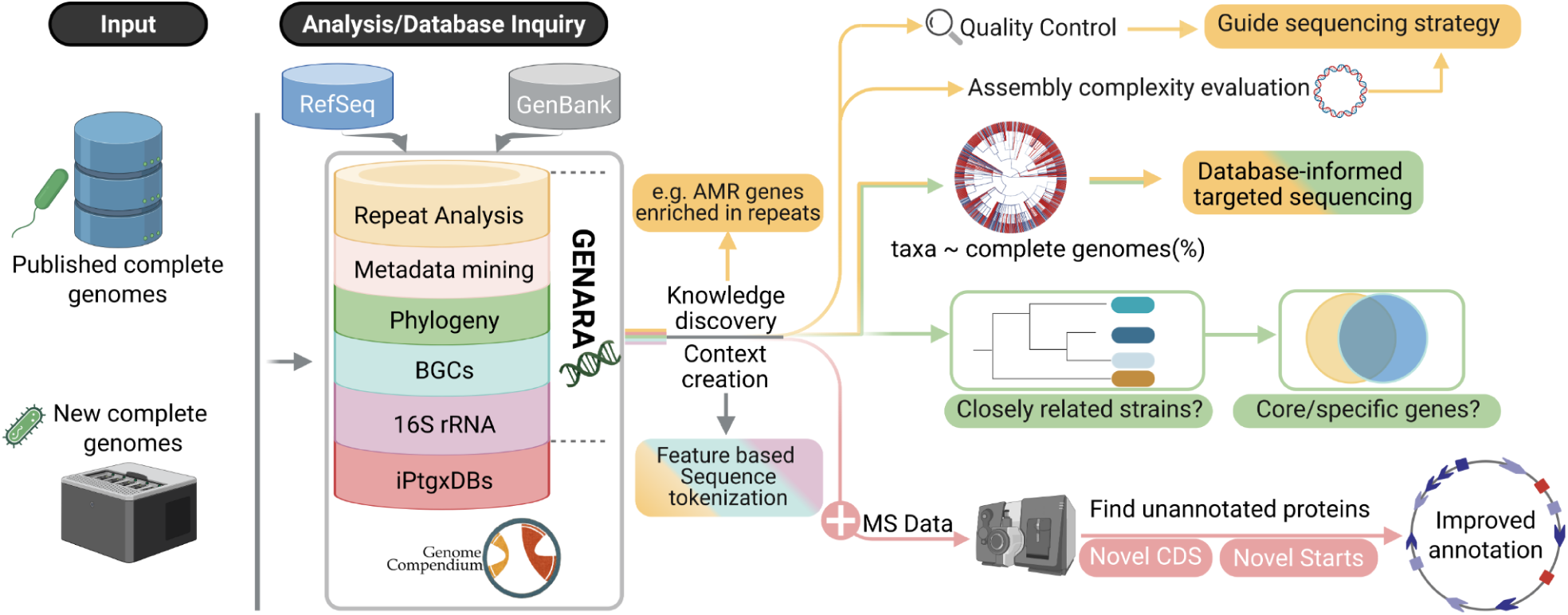

## Background

Microorganisms (MO), including prokaryotes, i.e., bacteria and archaea, are ubiquitous and perform many essential roles in diverse ecosystems. Cyanobacteria and selected other bacteria are primary producers that account for a substantial percentage of the oxygen and organic matter produced by photo- and chemosynthesis, which form the basis of most food webs(1). In soils, MO not only drive organic matter decomposition, nutrient cycling, water retention and carbon sequestration(2), but can also promote plant growth and suppress plant diseases(3). In fermented foods, bacteria, yeast and fungi drive metabolic processes that help to preserve food, enhance flavor and have probiotic effects(4). MO also have critical functions in bioremediation of contaminated soils(5) and as alternative, sustainable solutions to produce (and degrade) plastics(6), biofuels and chemicals(7). Consequently, there is increasing interest in exploring their biotechnological potential. Moreover, *Streptomyces* and other bacterial genera remain primary sources for discovering urgently needed antibiotics with novel modes of action(8, 9).

Scientists estimate that the vast majority of bacterial and archaeal species are not discovered yet, including many unculturable species, often referred to as microbial dark matter(10, 11). This implies that many beneficial microbes remain to be discovered, encoding diverse and potentially highly relevant functions. Large metagenomics studies support this view (12). A study of 27,000 metagenomes uncovered a large functional diversity among metagenome-assembled genomes (MAGs) that roughly doubled that encoded by reference genomes(13). Analyzing over 92,000 metagenomes, another study estimated that 10 to over 100 archaeal and bacterial phyla, respectively, and ∼80,000 bacterial genera are yet to be discovered(14). Importantly, metagenomics studies do not provide physical isolates, which are essential for transitioning from describing the composition of complex microbiomes by amplicon sequencing or metagenomics towards microbiome engineering(15). One promising strategy is to employ a consortium of functionally relevant strains to achieve a desired beneficial aspect, such as enhancing plant protection against pathogens(16). Studying synthetic communities (SynComs) enables researchers to decipher their interactions and to start identifying a mechanistic basis for these effects(17), ultimately informing the selection and application of optimal strain combinations. The prediction of particularly relevant strains from Syncoms data or even microbiome data represents a very promising approach(18).

Advances in next generation sequencing continue to shed light on the prokaryotic diversity and expand the number of sequenced genomes from strains of diverse microbiome isolates(19). Although short read sequencing has historically dominated, long read sequencing has gained popularity for its ability to generate complete prokaryotic genome assemblies. These represent an ideal basis for downstream functional genomics(20) and comparative genomics studies(21), and allow researchers to link varying phenotypes of closely related strains with genotype information(22). Moreover, complete genomes can be leveraged for a more accurate and comprehensive genome annotation by proteogenomics(23, 24), an important need of the community(25). The frequent repeat sequences in bacterial genomes still represent the main barrier to complete assembly. This applies in particular to short read-based assemblies, which are reported in the form of 10s-100s of contigs and thus lack a variable fraction of the genome including coding sequences (CDS) that can carry out important functions. In the case of *Pseudomonas aeruginosa* MPAO1, the missed CDS included genes with roles in immune evasion, antibiotics production, for competition with other bacteria and even essential genes(26). For very repeat-rich genomes, e.g. those of lactobacilli that can harbor hundreds of transposases, the fraction of missed/partial genes in short read assemblies can account for a few percent and include core genes(27). Importantly, CDS can also be missed in assemblies based on long read sequencing data from Pacific Biosciences (PacBio) and Oxford Nanopore Technologies (ONT), particularly in prokaryotic genomes harboring long, near-identical repeats well beyond 7kb in length(28). In 2018, we quantified the genome assembly complexity of 9,633 complete genomes from GenBank(29), and noted a substantial fraction of very complex to assemble genomes with long, near-identical repeats greater than 30 kbp (3.2%).

Our detailed analysis of repeat sequences in ∼60,000 complete prokaryotic genomes (RefSeq plus GenBank) revealed that complex prokaryotic genomes are much more prevalent than previously estimated. Integrating the repeat analysis with metadata for quality control enabled us to identify up to 6.3% of complete RefSeq genomes that contain potential assembly errors or may miss sequence information. We visualize the expanding taxonomic coverage over time and pinpoint opportunities for targeted sequencing to increase the representation of distinct species with complete genomes. Analyses of 16S ribosomal RNA (rRNA) intragenomic copy number and sequence divergence, biosynthetic gene cluster (BGC) content and the association of repeat sequences with specific AMR genes provide examples of relevant biological insights. We release the GenomeCompendium (https://genome-compendium.com/) as a dedicated database for this reference dataset. On top of integrating computations and metadata, we cross-reference external resources detailing strain availability, more comprehensive annotation and a genome-centric taxonomy, facilitating interactive online exploration of prokaryotic complete genomes. Users can construct interactive queries via the interface to generate custom plots, bulk download pre-computed genome reports, and -as a unique feature-analyse their own uploaded prokaryotic genome sequences with the GENARA (for <u>Gen</u>omic <u>A</u>nalysis of <u>R</u>epeats and <u>A</u>nnotations) web server (30). The GenomeCompendium is a versatile platform for data mining and use cases, allowing scientists to uncover interesting biology, both known and novel.

## Methods

### Download and processing of genome sequence data and metadata

Complete prokaryotic genome assemblies were downloaded from the National Center of Biotechnology Information’s (NCBI) RefSeq and GenBank (December 31, 2024)(31). From here on, we interchangeably use the terms genome and assembly. Relevant metadata including species, strain, biosample, assembly accession, assembly level, GenBank/RefSeq paired assembly information etc. (**Supplementary Table S1**), were parsed from the respective assembly summary text files for bacteria and archaea from both databases. Information about genome size, number and size of chromosomes/plasmids, number of predicted genes, CDS, sequencing technology used, reference-based or de novo assembly strategy etc. was parsed directly from the gbk/gbff files (where available) using Biopython v. 1.79(32). Taxonomic information (species, genus, family, order, class, phylum, kingdom, domain) was retrieved using python’s ete3 library (module: NCBITaxa) v 3.1.3(33) and from the Genome Taxonomy Database (GTDB)(34), see below. Throughout the manuscript, we use taxonomic names from the NCBI taxonomy database(35) unless noted otherwise. For some taxonomic ranks information was missing (e.g., 26 RefSeq assemblies had no genus entry); see **Supplementary Methods**. The overall workflow (**Supplementary Figure S2**) encompasses download, metadata extraction, repeat analysis (for details, see **Supplementary Figure S3**), additional computations/analyses and integration with selected useful public resources (see below). **Supplementary Table S1**, an extensive “summary table”, integrates the extracted metadata and information, results of computations and potential quality control issues, and is one potential starting point for data mining.

### Repeat analysis, repeat classification and visualization

Information about the number, length and type of genomic repeats (≥ 500 bp with a sequence identity ≥ 95%) was computed with a custom pipeline based on Nucmer v 3.1(36). Fasta sequences were extracted from the gbk/gbff files with Biopython and analyzed to identify and distinguish three types of repeat sequences: Interspersed repeats, tandem repeats and terminal inverted/directional repeats on linear contigs (see **Supplementary Figure S3**). The precise steps on how interspersed, tandem, terminal inverted and terminal duplicated repeats were identified, further processed and integrated into report files and summarized in **Supplementary Table S1** are detailed in the **Supplementary Methods**. The overall number of repeats and the maximum length of repeat pairs were used to classify genomes into three classes of genome assembly difficulty as described by Koren et al(28) (see Results). These values are also provided in **Supplementary Table S1**.

#### Repeat region content

To calculate the overall repeat region content, bedtools merge was used to combine overlapping repeats into single intervals. The length of all intervals was added up and divided by the genome length (**Table S1**).

#### Visualization

To visualize the number and length distributions of the three repeat types, swarm plots were created using the Python seaborn library. Terminal directional repeats were used mainly as quality control check (see below) and excluded from further analyses. The genomic location of the three repeat classes was also visualized with Circos (v.0.69-8,(37)). For interspersed repeats, the location of the respective repeat mates is shown. For visualization reasons, the plots show a maximum of 15,000 repeat pairs(∼2.1% of genomes exceeded that cut-off). Both plots are included in the individual genome assembly reports (PDF format).

### Proportion of complete genomes

NCBI RefSeq assemblies were filtered (both complete genomes and assemblies from the other three genome completeness levels) to include entries up to the end of 2024, and that contain a valid scientific name (genus and species tag, in some cases preceded by Candidatus; see **Supplementary Methods**). For each taxonomic rank, the proportion of genomes labeled as complete relative to the total number of assemblies in that rank was computed. An aggregated score was used as a proxy for genome assembly difficulty. At the species level, an average Koren score (an average assembly difficulty rating on a 1-3 scale) was computed where possible. For higher taxonomic ranks, the average of the values from the immediately lower taxonomic level was computed and used. Plots were generated using a number of additional R packages (see **Supplementary Methods**) to effectively represent taxonomic completeness (**Figure 1**) and assembly complexity across different ranks (**Figure 3**).

**Figure 1:**
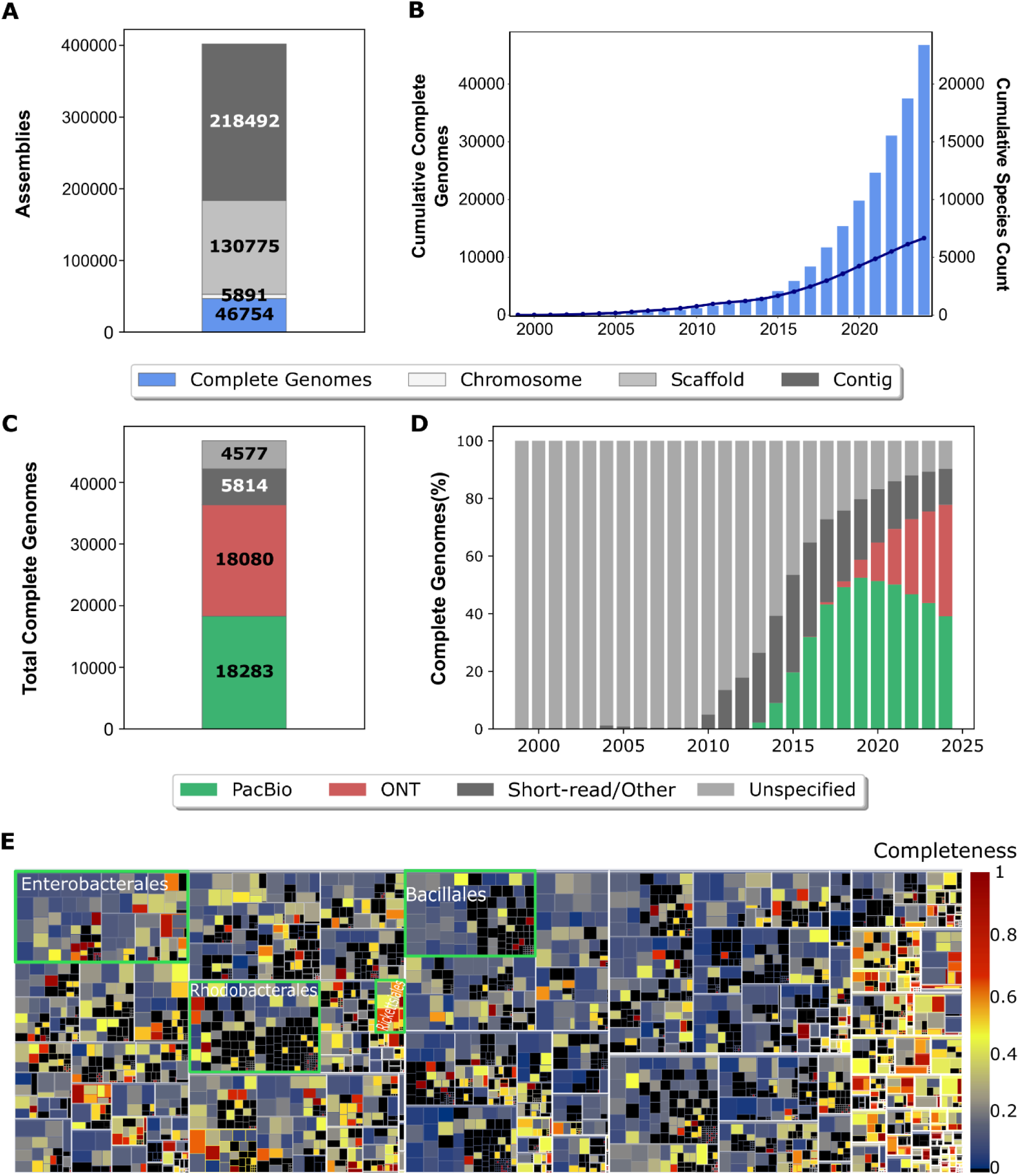
Prokaryotic genome assemblies at NCBI RefSeq. **A.** Stacked barplot of four classes of genome assembly completeness (see legend). **B.** Cumulative yearly increase of complete prokaryotic genome assemblies and distinct species (dark blue line). **C**. Stacked barplot with the number of complete genomes contributed by different sequencing technologies; PacBio (green), ONT (red), Short read/Other (predominantly short read seq. technologies like Illumina; dark gray) and unspecified (no metadata available, light gray). **D**. Cumulative increase of complete prokaryotic genomes over time and per technology category. Note: for C and D, PacBio+ Illumina assemblies are counted as PacBio; ONT + Illumina as ONT, PacBio + ONT (60 cases) as ONT. **E**. Treemap at the genus level highlighting the proportion of complete RefSeq genomes (completeness). Box sizes are based on the logarithm of the number of genomes, genera with no complete genomes are shown in black (for an interactive version of this Figure, see https://genome-compendium.com/). Selected higher taxonomic ranks (order) are labeled by green boxes and shown with white headers.

### Quality Assessment

Metadata extracted from the respective RefSeq/GenBank files was used to flag assemblies that relied on a reference-based assembly strategy (for all or a subset of the replicons), as well as genomes that were assembled using only data from short read technologies (Illumina, 454, Sanger, ABI, iontorrent, dnbseq, SOLiD) but where our repeat analysis identified “long” repeats > 7000 bp (using Koren’s definition)(28). We also flagged assemblies that contain a linear contig for which we detected terminal directional repeats, which could indicate a misassembly, e.g. a missing circularisation step for a chromosome or plasmid. Visual inspection of Circos plots provided in the report files can help users cross-check assemblies and identify potential issues, as assembly algorithms still struggle with duplicated contigs and repeats shared between plasmids and chromosomes.

### GTDB Taxonomy

To obtain an additional genome-based taxonomic classification, the Genome Taxonomy Database (GTDB) was used(34). Most assemblies were included in release R226 and the respective classification was transferred using their NCBI assembly identifiers. Assemblies not present in R226 were classified from FASTA sequences using the GTDB-Tk classify_wf workflow (v2.4.0, default parameters)(38). The differences of NCBI and GTDB classifications are summarized in **Supplementary Figure S8**.

### Enrichment analyses

Hypergeometric tests were performed on RefSeq assemblies with assigned genus or family names (46,728), excluding partial or unresolved 16S rRNA annotations (8 cases) and using the Holm multiple testing correction(39). We assessed taxonomic enrichments in genomes for several features: repeat frequency/type (absence, terminal inverted or directional, long, or abundant repeats), assembly type (reference-based or mixed), Koren class II genomes, 16S rRNA copy number/sequence identity threshold, and high overall BGC counts. Note: for enrichment analyses at the rank species, we considered the species_defined column (see **Supplementary Methods**). For each taxonomic rank, the observed number of assemblies with a given feature was compared to the total available number of assemblies. Results are reported as absolute counts, percentages relative to the total assemblies per taxonomic rank, and adjusted p-values in **Supplementary Table S2**.

#### AMR gene enrichment

All CDS of RefSeq genomes with a valid species and genus name were searched for the broader gene categories “antimicrobial resistance”, “stress” and “virulence related genes” using AMRFinderPlus v.4.2.7 (40). rRNA features were extracted from the annotated type field, transposases and ribosomal proteins from the annotated product field. Only genera with at least 3 species, each with at least 2 genomes and at least 2 features per genome, were considered for this enrichment analysis. Enrichment within identified repeats was performed using a Mantel-Haenszel test stratified by species on the log odds ratio (OR), complemented by a random-effects model on the per-stratum estimates. Features were required to overlap a repeat by at least 70% and were considered significant when the absolute log_2_OR>2 and both Benjamini-Hochberg corrected Mantel-Haenszel and random-effects p-values were below 0.05. For more detail, see **Supplementary Methods**.

### rRNA gene analyses

The 5S, 16S and 23S rRNA gene coordinates were parsed from the RefSeq annotation and their respective lengths computed (**Supplementary Table S1**). Additionally, the 16S rRNA fasta sequences were extracted using Biopython (v1.81(32)). Eight assemblies with reported incomplete 16S rRNA genes and 26 assemblies that lack a genus assignment and information about their 16S rRNA gene(s) were not considered in this analysis of 46,720 genomes. For assemblies with at least two 16S rRNA copies, global optimal pairwise alignments between all 16S rRNA sequences were generated using VSEARCH(41) using the option –allpairs_global. Percent identity was defined as the edit distance, excluding terminal gaps (id2 in VSEARCH); see section 16S rRNA divergence in **Supplementary Methods**. Results are also available in the report files. Where available, the copy number of 5S, 16S and 23S rRNA genes is listed (**Supplementary Table S1**). Finally, an analysis of rRNA operons and their location in specific repeat types was carried out (for details, see **Supplementary Methods**).

### Analysis of biosynthetic gene clusters

To detect putative biosynthetic gene clusters (BGCs), we ran antiSMASH v.8.0.1(42) on the FASTA sequences of all assemblies using Prodigal as gene finder. The protoclusters that are identified consist of a core region of genes with its surrounding neighborhood and are predicted to be involved in the synthesis of a single product type. Protoclusters thus represent regions of interest that may encode a BGC. The option knownclusterblast was selected to compare identified protoclusters to known clusters from the Minimum Information about a Biosynthetic Gene cluster (MiBiG) database(43). Predicted BGC types were extracted from the summary GenBank file, looking for all “protocluster” features. From the summary text files provided for each region of biosynthetic activity, the BGC identifiers of significant hits to the MiBiG repository (as defined by antiSMASH) were extracted. For each hit, we also report the number of genes in the region that match genes in the associated MiBiG cluster. The number and types of predicted BCGs per region, the BGC identifiers with their respective gene hits, and total gene count per region were added to both **Supplementary Table S1** and report files. Finally, the RefSeq genomes were analyzed to identify genera that preferentially harbor many protoclusters, outliers within a specific genus, and co-occurrences of particular protoclusters.

#### Data mining and visualization

Results from the analyses described above were incorporated in the summary table (**Supplementary Table S1**). Figures were generated using matplotlib, seaborn and plotly. Statistical analyses including hypergeometric tests, lowess regression, PCA, percentiles were carried out with python’s scipy, statsmodels and sklearn libraries. Taxonomic trees (Figures 5, S11) were constructed with NCBI’s taxonomic identifiers using the R package ape (version 5.8) and plotted with ggtree (version 3.12.0). The trees in Figures S9B, S11B were plotted with R package metacoder (version 0.3.7)(44).

### Crosslinks to selected resources

#### Link to proGenomes

This very useful resource provides an inventory of close to 2 million genome assemblies (complete and fragmented) that have been consistently and uniformly annotated(19). Where possible, we provide a link to the respective proGenomes entry (**Supplementary Table S1**).

#### StrainSelect

we mined the StrainSelect dataset(45) to link a subset of the complete genomes to a Bioresource center (BRC) from which they can be purchased. This info is available in **Supplementary Table S1** and in the individual report files; an overview of how many strains are available per BRC is shown in **Supplementary FigureS14**.

#### Exploring increase of prokaryotic genomes over time

NIH genome sequencing cost data were downloaded from www.genome.gov/about-genomics/fact-sheets/DNA-Sequencing-Costs-Data and used to identify specific time periods during which reductions in sequencing costs corresponded with accelerated growth in the number of completely sequenced genomes.

### GenomeCompendium public web server

#### Infrastructure, results download and online data table

The results of the repeat analyses, of the quality control assessment, of additional computations, metadata and crosslinks to selected other resources (where available) are publicly available via our cloud-based web server at https://genome-compendium.com/. For each complete genome, we provide a detailed report in PDF format that includes both tabular information and several figures. Information from **Supplementary Table S1** is made available as an online asynchronous access PostgreSQL database. Through the online interface, users can construct and apply multiple queries to filter the data table and generate plots for any selected column(s) of interest. Additionally, they can download input sequences, antiSMASH result files, and raw repeat identification files. The website also provides interactive summary diagrams that users can explore for further insights. To download the available resource files automatically, dedicated URLs can be used (e.g. genome-compendium.com/Report/GCF_038627105). Complete usage instructions are provided in the documentation available on the web server. The platform was developed as a Golem Rshiny package (46) and relies on additional R packages (**see Supplementary Methods**).

#### GENARA webtool

Users can upload any complete prokaryotic genome sequence and run it through our online GENARA analysis tool to perform the following analyses: repeat identification and summarization, GTDB taxonomy classification, 16S rRNA gene identification plus sequence similarity computation and integration of antiSMASH predictions. Briefly, GENARA takes FASTA or GBK files as input and runs a series of orchestrated Podman containers for each analysis step. For each user, a background process is started, allowing them to close their browser without losing progress. If desired, users can be notified by email when their analysis is completed. The analysis results are retained on the website for a maximum of five days. Detailed information on using GENARA is available online. The containers used to run the GENARA analysis tool are also available as prebuilt versions (10.5281/zenodo.18247865) as well as built codes with instructions (https://github.com/mgbteam/GENARA).

### Proteogenomics search databases as a basis to improve genome annotations

For each completely sequenced prokaryote in NCBI RefSeq (46,754; Dec. 2024), we created a custom integrated proteogenomics database (iPtgxDB) as described(23), allowing researchers to improve the genome annotation of their favorite organism(s) using proteomics data. Here, we deliberately integrated only a few, selected annotation/prediction sources in order to create relatively small iPtgxDBs comparable in size to the RefSeq search databases, which offers some benefits(24, 47). These iPtgxDBs integrate annotations from NCBI’s PGAP(48) (included in the downloaded GenBank files), from the *ab initio* predictor Prodigal (version 2.6.3)(49), and predictions of small ORFs (smORFs) encoding proteins below 100 amino acids based on GMSC-mapper, querying the global microbial smORFs catalog (GMSC)(50). Prodigal was run with the closed ends option enabled and masking stretches of Ns (-c and -m flags), while GMSC-mapper was used with default options. For most genomes, we relied on bacterial and archaeal translation code 11, except for 783 assemblies (12 genera from the phylum Mycoplasmotata, and 73 assemblies from the genus *Providencia*), which use translation code 4, and one assembly that uses translation code 25 (*Candidatus Absconditicoccus praedator*). Any of the 46,754 iPtgxDBs can be downloaded from our public GenomeCompendium web server (https://genome-compendium.com/). For researchers interested in capturing the full coding potential, larger standard iPtgxDBs can readily be created (please see the dedicated website and documentation of this separate web server: https://iptgxdb.expasy.org).

## Results

### Complete prokaryotic genomes - a small but growing fraction of assemblies

The NCBI classifies genome assemblies into four categories of completeness and data quality: complete genomes (the highest quality level), chromosomes, scaffolds and contigs, the lowest level of assembly completeness(31). Among 401,912 prokaryotic genome assemblies available from RefSeq (end December 2024), 46,754 (11.6%) were classified as complete genomes, 1.5% as chromosomes, 32,5% as scaffolds and 54.4 % as contigs (**Figure 1A**). The cumulative increase of complete genomes (**Supplementary Figure S1**) and distinct species exhibited an average annual increase of 17% ± 4.6% in newly identified species since 2015 (**Figure 1B**; dark blue line; scientific names are from the NCBI taxonomy unless specified otherwise). When separated by sequencing technology, assemblies based on 3rd generation long reads from PacBio (green) and ONT (red) cumulatively accounted for 39.1% and 38.7% of complete genomes in 2024 (**Figure 1C**), respectively. The first PacBio-based genome assemblies appeared in 2013, followed by ONT assemblies in 2015 (**Figure 1D**), with the latter surpassing PacBio in volume in 2022 (**Figure S1**). While complete genomes assembled from short read data alone -requiring time-consuming manual curation-dominated until 2012 (**Figure S1B**), their relative yearly proportion decreased thereafter, accounting for roughly 12.7% in 2024.

Compared to 9,633 prokaryotic GenBank genomes analyzed in 2018 (9,331 bacteria, 293 archaea;(29)), the RefSeq2024 dataset (complete genomes of 46,143 bacteria and 611 archaea 1.3%; **Table 1**), represents a near 5-fold increase mainly driven by bacteria. GenBank2024 (∼2,45 million) has 6 times as many assemblies as RefSeq, with larger percentages in the categories scaffold and especially contigs (**Figure S1D-E**). As the criteria for inclusion in RefSeq are more stringent than for GenBank, we only considered 13,224 additional complete GenBank genomes, for overall 59,978 complete assemblies. Major fractions of these additional genomes stem from i) large multi-isolate pathogen sequencing projects, ii) metagenome projects and iii) genomes of undefined genus or unverified source origin (**Figure S1F**). Most of our analyses focus on the more stringent RefSeq dataset (**Supplementary Table S1** also lists results of selected analyses for complete GenBank assemblies). A first analysis of the RefSeq dataset revealed that the proportion of complete genomes among the taxonomic ranks covered overall differs substantially. This is shown in **Figure 1E** at the genus level with selected higher ranks labeled (here for the orders Enterobacterales, Bacillales and Rhodobacterales, green boxes), illustrating the variable amount of black squares that show taxa (here genera) lacking complete genomes (see Methods). This directly implies that targeted sequencing efforts for genera where no complete genome but only fragmented assemblies exist could increase the overall number of distinct species covered (**Figure 1B**). A list of genera with no or fewer than 20% complete genomes is shown in a zoomable Figure on the web server website.

**Table 1.** Prevalence of repeat types in bacterial and archaeal genomes.

| Category | Bacteria |  | Archaea |  |
| --- | --- | --- | --- | --- |
|  | number | % | number | % |
| Analyzed prokaryotic genomes | 46,143 | 100 | 611 | 100 |
| Genomes without repeats* | 387 | 0.8 | 54 | 8.8 |
| Genomes with interspersed repeats | 45,552 | 98.7 | 540 | 88.4 |
| Genomes with tandem repeats | 41,735 | 90.4 | 339 | 55.5 |
| Genomes with terminal inverted repeats | 740 | 1.6 | 1 | 0.2 |

\*selection criterium: repeat sequence of $\geq 500$ bp with $\geq 95\%$ nucleotide identity. Note: the repeat classes (interspersed, tandem, terminal inverted) can overlap to some degree.

### Analysis of repeat sequences in prokaryotic genomes

The length and number of repeat sequences are parameters that profoundly affect *de novo* genome assembly. Near-identical repeats (≥ 500 bp, sequence identity ≥ 95%) were thus used to define 3 ‘Koren’ classes of assembly difficulty for prokaryotic genomes(28): Class I genomes that are straightforward to assemble (longest repeat is the entire ∼7 kb rDNA operon, overall less than 100 repeats), class II genomes (longest repeat is the rDNA operon, but more than 100 repeats), and class III genomes, which can harbor repeats substantially longer than the rDNA operon and that can require major effort to assemble a contiguous, complete genome. Applying this classification to 9,633 complete GenBank genomes (36.2% class III) in 2018, we provided a first global estimate of very difficult to assemble class III genomes with repeats of 30kb or longer (3.2%)(29).

We here substantially extend the repeat analysis of prokaryotic genomes and integrate relevant metadata to create a basis for data mining and quality control. We distinguish three different types of repeats: interspersed repeats (two or more repeat units dispersed across the genome), tandem repeats (two or more repeat units directly adjacent to each other) and terminal inverted repeats (TIR) that can be found at the end of linear bacterial chromosomes(51) and plasmids(52) (**Table 1**, **Figure 2A**). When visualizing the repeats, we also identified terminal directional repeats on linear contigs, some of which may represent assemblies with a potential quality control (QC) issue, i.e., a missed circularization step for contigs that might represent circular plasmids or chromosomes (**Figure 2A**). An overview of our workflow covering metadata extraction, repeat classification, additional analyses and cross-references is shown in **Supplementary Figure S2**, a detailed description of the steps of the algorithm to identify the three repeat types is provided in **Supplementary Figure S3.** To simplify the visualization, we combine repeats into repeat regions if smaller repeat(s) are completely contained within a longer repeat, or if two repeats overlap by more than 95% of the smaller repeat (**Figure 2B**). We also calculate the repeat region content, i.e., the fraction of a genome that is covered by repetitive sequences (**Supplementary Figure S4**). Using our cut-offs, the median repeat region content in RefSeq2024 is 3.2%. The top repeat region content value for RefSeq genomes is 66% (a Spiroplasma endosymbiont of the beetle *Clivina fossor*). A value of 54% was observed for *Spiroplasma ixodetis, a* cell wall-less, genome-reduced (∼2.2 Mb), gram positive bacterium of the class Mollicutes. It is a symbiont of ticks and other arthropods where it can lead to male killing of its hosts(53). The genome harbors many copies of transposase families (IS1595, IS5 and IS3, ∼950 to 1350 bp) that are located on plasmid 2 and the chromosome; overall, they account for more than 450 repeats (**Figure 2D**).

**Figure 2.**
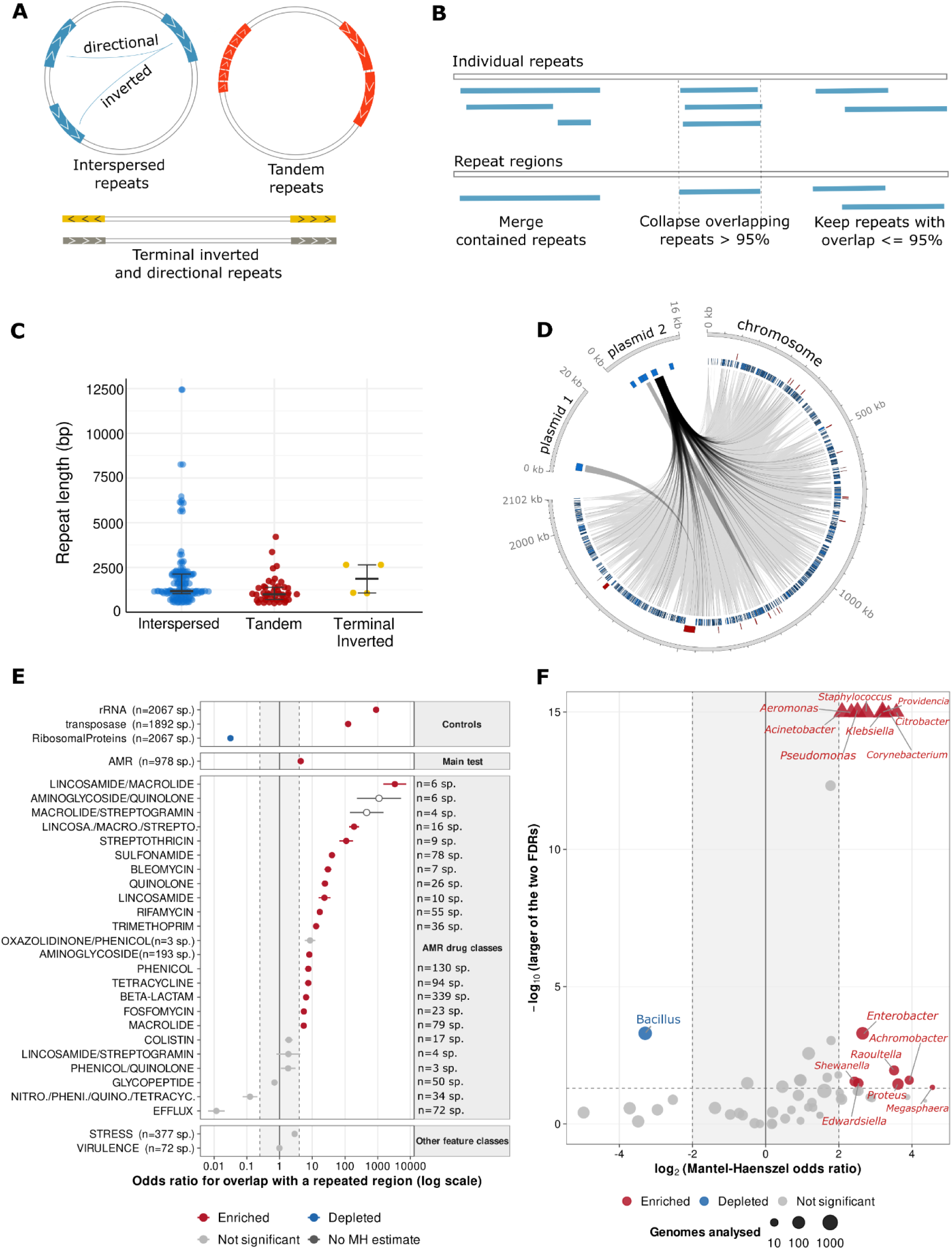
Overview and visualization of repeat types, and enrichment of gene classes in repeats. **A.** Schematic overview of the three types of repeat classes that we differentiate: interspersed repeats (either directional or inverted, blue), tandem repeats (two or more copies, red) and terminal inverted (golden) or terminal directional (gray) repeats. **B.** Combination of repeats into repeat regions (also see **Supplementary Figure S4**). **C.** The occurrence and length of repeats in the three classes is shown in swarm plots; they are part of the downloadable reports. Here, the cyanobacterium *Nostoc sp.* is shown (GCF_002949735.1), which harbors repeats of all three classes; the repeat length values indicate that this is a class III genome (repeats > 7 kb). **D.** Circos plot of the genome of *Spiroplasma ixodeti*s with one of the top repeat region content values (0.54) in RefSeq (GCF_027923845.1; one chromosome (2.1 Mb), two plasmids (20.1, 16.4 kb)). The majority of the overall 956 repeats regions are interspersed (934; blue boxes), the rest are tandem repeats (22; red boxes). **E.** Mantel-Haenszel (MH) odds ratios (ORs) with 95% confidence intervals for feature-class overlap with repeat regions, stratified by the associated species. The gray band indicates an OR of 0.25-4; coloured points outside this range meet both FDR significance thresholds (<0.05) and are classified as enriched (red) or depleted (blue). Open symbols indicate no MH estimate; the random-effects estimate is shown instead. **F.** Genus-level enrichment of AMR genes in repeat regions, with species as strata. The y-axis shows the more conservative of the BH-corrected MH and random-effect p-values. Genera outside OR 0.25-4 and meeting both significance thresholds are again classified as enriched or depleted. Triangles indicate capped y-values (see **Supplementary Table S2** for the precise values).

With the repeat cut-offs used (see Methods), we identified 441 RefSeq assemblies without repeats (<1%); they are more common in archaea (8.8%) than in bacteria (below 1%; **Table 1**). An enrichment analysis (see Methods) revealed several genera that were over-represented including *Buchnera* (61 of 79 assemblies without repeat), *Blattabacterium* (28/43), and the three *Candidatus* genera *Karelsulcia* (56/59), *Blochmaniella* (21/26) and *Carsonella* (15/17) (**Supplementary Table S2**). These bacteria are all obligate endosymbionts of insects that have co-evolved with their hosts and have undergone substantial genome reduction. A summary of the number of repeat pairs per type and their lengths are shown in so-called swarm plots **Figure 2C**. Together with a Circos plot that visualizes the genomic locations of the three repeat classes (**Figure 2D**), they are provided in a report file for each assembly.

Interspersed and tandem repeats are the most frequent types and found in over 90% of bacterial genomes (**Table 1**). Interspersed repeats account for over 92.2% of the roughly 3.1 million repeats recorded in all genomes, while roughly 237,000 tandem repeats (7.7%) were observed, which are less frequent in archaea. The length of the longest repeat per repeat type and the range of occurrences are detailed in the **Supplementary Results** and in **Supplementary Figure S5.** Assemblies with TIRs (1.6% of bacteria) were highly enriched in the genera *Streptomyces* (625 of 1253) and *Borrelia* (41 of 78; **Supplementary Table S2**). In *Streptomyces*, TIRs were predominantly found at the end of linear chromosomes(51), in *Borrelia* predominantly on linear plasmids(52); yet, a small fraction of assemblies from these genera also harbored TIRs on both plasmids and chromosomes. Finally, terminal directional repeats were smaller (up to 87.8 kb) and found in 191 genomes.

Finally, we explored whether specific gene categories and classes were enriched within repeat regions (see Methods and **Figure 2E, F**). As expected, we observed a strong positive enrichment for transposases (log_2_OR>6.95, FDR<0.01) and rRNA genes (log_2_OR>9.8, FDR<0.01) that are associated with repeats(54, 55). In contrast, a significant negative enrichment was observed for ribosomal proteins (log_2_OR<-5.05, FDR<0.01)(56) (**Supplementary Table S2**). Previous work has shown that transposition of mobile genetic elements can generate duplications of antimicrobial resistance genes (AMR), and that such duplications are favoured under antibiotic selection(57). Within our dataset (**see Supplementary Methods**), AMR-related genes overall were significantly enriched in specific species (**Figure 2E**). Notably, we found certain gene subclasses to be overrepresented within repeat regions, such as sulfonamide resistance genes (log_2_OR>5.3, FDR<0.01) and quinolone resistance genes (log_2_OR>4.6, FDR<0.01) (**Figure 2E**), both of which have previously been reported to be associated with integrons, which contribute to multidrug resistance in *E. coli* strains(58, 59). Streptothricin resistance genes were also strongly enriched within repeated regions (log_2_OR>6.7, FDR<0.01).

Evaluating the association between repeat regions and AMR genes across different genera, we found a clear separation between genera in which AMR genes tend to occur within repeated regions and those in which they do not (**Figure 2F** and **Supplementary Table S2**). *Bacillus* genomes had one of the lowest association rates between repeat regions and AMR genes: only 23 of 10,693 AMR genes analyzed for this genus were located within repeat regions. In contrast, *Citrobacter* (log_2_OR>3.57, FDRs<0.01) and *Proteus* (log_2_OR>3.61, FDRs<0.01) each have over 20% of their AMR genes associated with repeat regions. Among the analyzed RefSeq genomes, *Empedobacter* had over 40% of its AMR-associated genes located within repeat regions; due to the small sample size (24 out of 59 genes across 6 genomes) this however was not statistically significant. With the exception of *Enterococcus*, all ESKAPE pathogen related genera and specific species showed an enrichment of AMR-associated genes within repeated regions (log_2_OR>2.08, FDRs<5×10-4), with *Klebsiella pneumoniae* presenting the highest overall proportion of repeat-associated AMR genes (17%), **Supplementary Figure S6**.

### Repeats affect genome assembly complexity

The RefSeq2024 dataset contains 2,724 genomes whose longest repeat is ≥ 30kb (5.8%), i.e., a large increase compared to the 3.2% reported earlier(29). An enrichment analysis identified the genera *Streptomyces*, *Leptospira*, *Pseudomonas* and *Acinetobacter* to be over-represented among these very complex genomes (**Table S2**). Advances in long read sequencing technologies thus helped to increase the fraction of very complex genomes, a trend we visualize in **Figure 3**. Mining the available metadata provided evidence that over the years, genomes with increasingly longer repeats and higher overall number of repeats and repeat region content (**Figure 3A**) could be fully resolved. The increasing percentage of difficult to assemble class III genomes that were submitted to RefSeq (cumulatively) **(Supplementary Figure S7)** also support this conclusion. For genomes that were submitted to RefSeq after our analysis in 2018, 44.6% belong to class III. Compared to 36.2% of the assemblies submitted up to that time point (**Supplementary Table S3**), this is a statistically highly significant enrichment (P-value < 3.0e-47). Using a double logarithmic scale, all assemblies can be visualized, revealing that genomes can harbor well over 1000 repeats (**Figure 3B**; top entry: *Endozoicomonas* sp. with 1398 repeats (NZ_CP092417.1; **Table S1**), and have maximal repeat lengths over 2 Mb (top 2 entries: *Streptomyces* sp strains NBC_01613 and NBC_00496 with TIRs of 2,415,238 bp (GCF_045209205.1) and 2,372,486 bp (GCF_036013585.1), **Table S1**). An integrated analysis of genera further stratified by their mean numerical Koren class score (see Methods) and the fraction of complete genomes among all RefSeq genomes is shown in **Figure 3C**. Strains from the prominent genus *Corynebacterium* (upper left corner) are overall relatively easy to assemble (mean Koren class <1.25); the fraction of complete genomes of close to 30% supports that. Members of the genera *Pseudomonas* and *Streptomyces* are on average more difficult to assemble (mean Koren class just below and above 2.25, respectively, with a complete genome fraction slightly above 10%). Genera with overall very difficult to assemble genomes (mean Koren class >=2.5) for which the overall fraction of complete genomes is below 20% (example genera: *Vibrio* and *Pedobacter* in the light-blue labeled rectangle) are shown as a taxonomic tree in **Figure 3D**. They represent groups where long read sequencing, ideally combined with efforts to extract intact high molecular weight genomic DNA, is likely required to generate additional complete genome assemblies.

**Figure 3.**
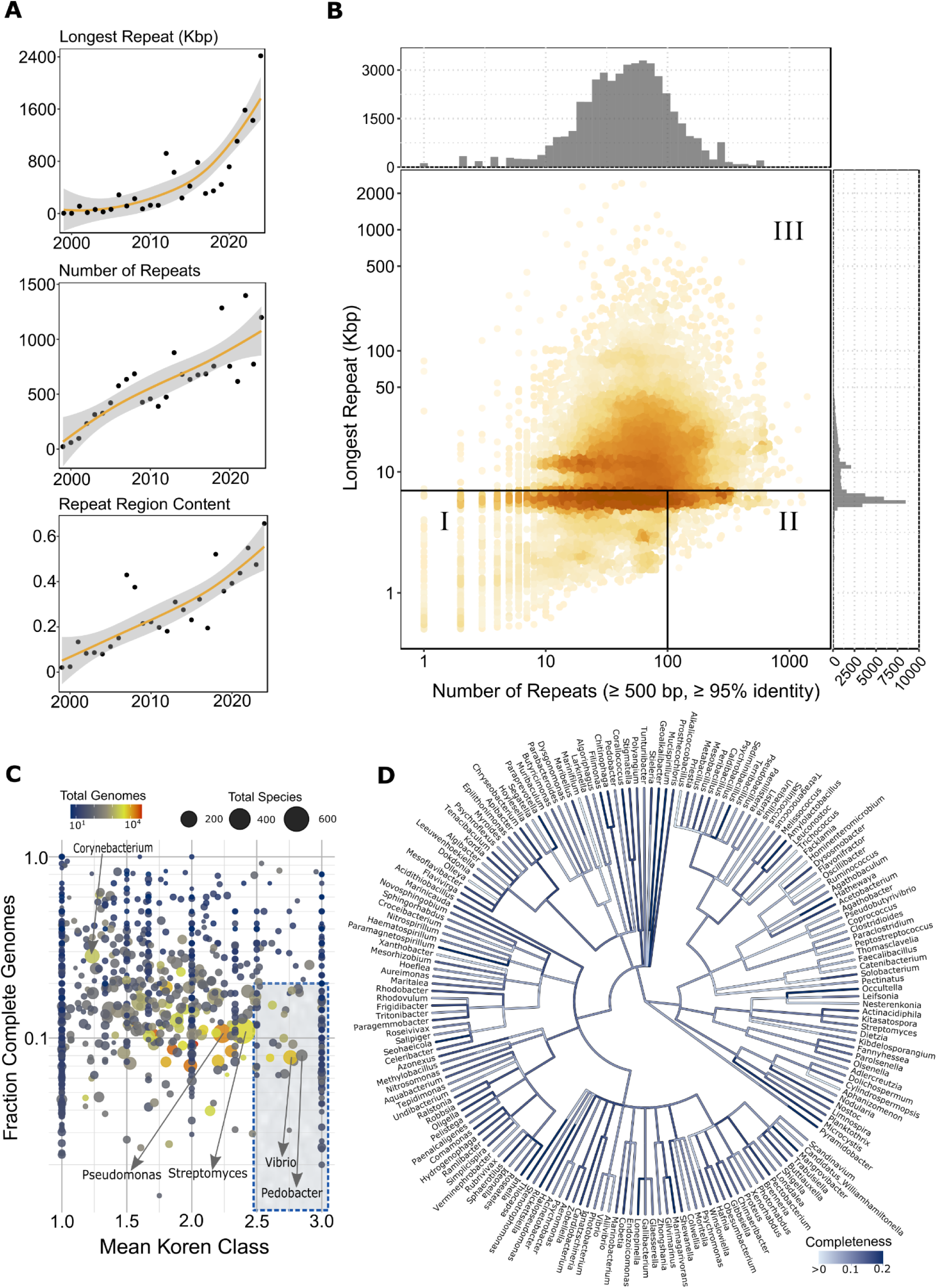
Resolving increasingly complex genomes over time. **A**. Advances in sequencing technologies allow us over time (x-axis)to resolve increasingly complex genomes with respect to their longest repeat length, total number of repeats and higher repeat region content (y-axes). The orange line shows the LOWESS regression line, while the gray cone represents a 95% confidence interval. **B.** A log log version of the original Koren plot captures the full range of genome assembly complexity for all RefSeq genomes. A heat map and histograms on top and to the right show how the genomes are distributed in terms of number of repeats and longest repeat. **C**. The fraction of complete genomes for different genera (with at least 5 assemblies) versus mean numerical Koren class score is shown, colored by the number of total genomes. A square of genera with particularly difficult to assemble genomes is highlighted and selected genera shown as examples (*Vibrio*, *Pedobacter*). **D.** Taxonomic tree of the genera contained within the blue-dashed square in Figure 3C for which less than 20% of the strains have a complete genome (completeness). Branch colors can change when higher taxonomic ranks get aggregated and the respective values change.

The hypothesis that the assembly complexity of not yet sequenced members of a genus can be estimated from the members with complete genomes available can be justified: Among genera with more than 5 complete assemblies, 60% of genera were dominated by a single Koren assembly difficulty class (more than two thirds of assemblies belonged to the same class). A logistic mixed-effects model with genus as a random effect yielded an intraclass correlation (ICC) of 0.37, indicating a moderate tendency for difficult to assemble genomes (class 3) to co-occur within the same genus. On the website, we show a detailed taxonomic tree for genera represented among RefSeq genomes for which either no complete genome is available, or the fraction of complete genomes is below 20%. This again enables interested users to prioritize sequencing efforts in areas that will most effectively improve complete genome representation, an important use case of the GenomeCompendium.

### Quality aspects of RefSeq assemblies

One of our key objectives was to mine the available metadata for potential quality issues and to flag such assemblies. One obvious category to check concerned the subset of genomes where the metadata indicated that they were assembled using a reference-based approach (**Figure 4A**), either for all, or for a subset of the contigs in an assembly (1,351 cases; **Table S1**). An enrichment analysis uncovered genera that contain several pathogenic species to be enriched, including *Mycobacterium*, *Corynebacterium*, *Helicobacter* and *Xyllela* (**Table S2**). A *de novo* assembly is considered state of the art, in particular when long reads are available to correctly assemble even complex, repeat-rich genomes. It thus makes sense to flag assemblies that have used a reference-based assembly strategy, as they can miss sequence information that is not present in the reference strain. A reference-based assembly strategy was still employed for a small fraction up to 2024, although the percentage of affected genomes has been gradually decreasing from greater than 10% in 2014 to below 2% in 2024 (**Figure 4B**). Moreover, another subgroup of assemblies with potential errors are those that were assembled exclusively based on short read data, but for which our repeat analysis identified repeat pairs over 7000 bp in length. We identified 1,678 such cases (**Figure 4A**) and provide a histogram of the size of their longest repeats, which range from 7 to 921 kb (**Figure 4C**), thereby underlining the potential for mis-assemblies among these. Also shown are 191 genomes with terminal directional repeats (yellow circle), which may represent missed circularizations of chromosomes or plasmids (**Figure 4A**). The visualization of repeat pairs in the Circos plots that are part of the report file for each strain (see **Figure 2D**) is also useful to inspect assemblies for potential problems. Overall, the data imply that even for a curated database, some of the entries, i.e., up to 2,934 cases (6.3 %) for this release may have a potential quality issue. For most of them, a combined use of long reads and a de novo assembly strategy is expected to resolve the issue.

**Figure 4.**
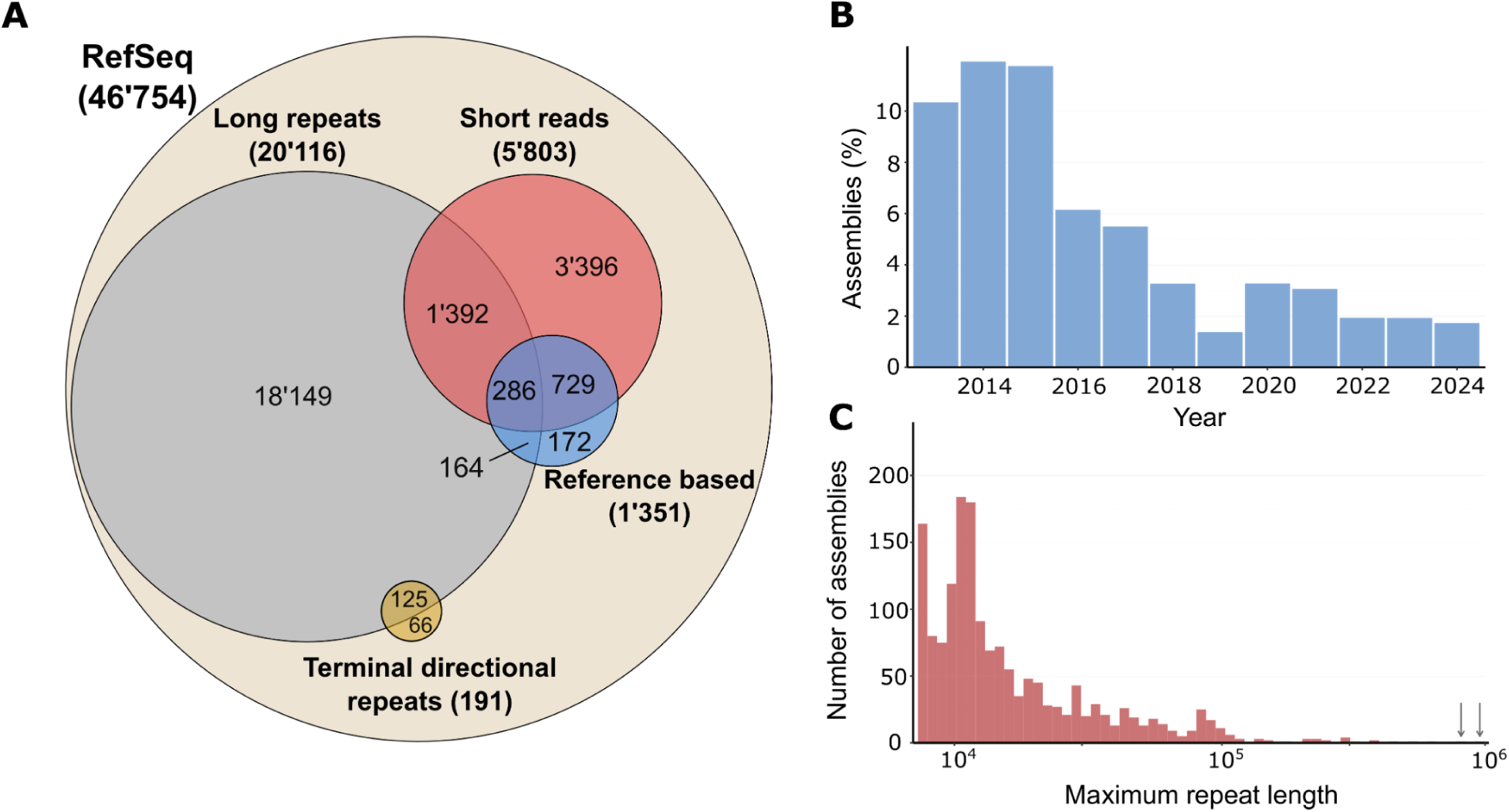
RefSeq genomes with a potential QC issue. **A**. A Venn diagram highlights the subset of the RefSeq genomes for which we identified potential QC issues. This includes 191 genomes with terminal directional repeats. Overall, 1,351 genomes were assembled with a reference-based assembly strategy (blue) and thus can miss sequence information. **B**. Percentage of genomes deposited in RefSeq that used a reference-based assembly strategy over the last 12 years. **C**. Histogram of 1,678 genomes (red) that harbor long repeats greater than 7 kb but are assembled only from short read data. Their maximum repeat length is shown. The two top entries have repeats of 783 kb and 921kb (grey arrows).

### Quantifying the increasing taxonomic coverage of RefSeq assemblies

We next assessed the taxonomic diversity of the growing collection of completely sequenced prokaryotes. As a first step, we retrieved or computed (see Methods) the taxonomy assignment from the genome taxonomy database (GTDB). It takes a different approach than the NCBI and provides a “phylogenetically consistent and rank normalized genome-based taxonomy for prokaryotes”(34). When comparing NCBI RefSeq and GTDB classifications, most differences were observed at the ranks species and order, followed by family and then class (**Supplementary Figure S8**). Notably, for a subset of 5,645 RefSeq assemblies, our parsing did not detect a valid scientific name, but only a genus name without resolution of the species (see **Supplementary Methods**). For 41,2% of these entries (2328/5645), the GTDB taxonomy does provide a specific species name/identifier and thus a higher resolution (**Table S1**). To assist researchers in selecting the taxonomy most appropriate for their needs, we provide both classifications for RefSeq and GenBank assemblies (**Table S1**).

For the categories family, genus and species, the number of new distinct taxonomic entries added per year from 1999 to 2024 is shown in **Figure 5A**, for the higher order ranks in **Supplementary Figure 9A**. While the linear fit becomes better with more data points, a trend can be identified that the number of novel higher order taxonomic ranks (phylum, order, class) added per year is declining (**Supplementary Figure 9A**). In contrast, for the ranks family, genus and species, there is a yearly overall increase: roughly 500-600 distinct new species were added per year since 2020 and 130-140 new genera, respectively (**Figure 5A**). Among the species with most assemblies in the dataset, *Escherichia coli* ranked first (3,726 genomes), followed by *Klebsiella pneumoniae*, *Salmonella enterica*, *Staphylococcus aureus* and *Pseudomonas aeruginosa* (**Supplementary Figure S10**), i.e., some of the infamous ESKAPE pathogens that include various multi-drug resistant bacteria for which novel therapies are urgently needed(60). However, *Bacillus velezensis* (410 genomes), a species that plays important roles as plant growth promoting bacterium (PGPB) and that can provide protection against phytopathogens(61), and *Bacillus subtilis* (343 genomes), were also among the top 20 species (**Figure S10**).

**Figure 5.**
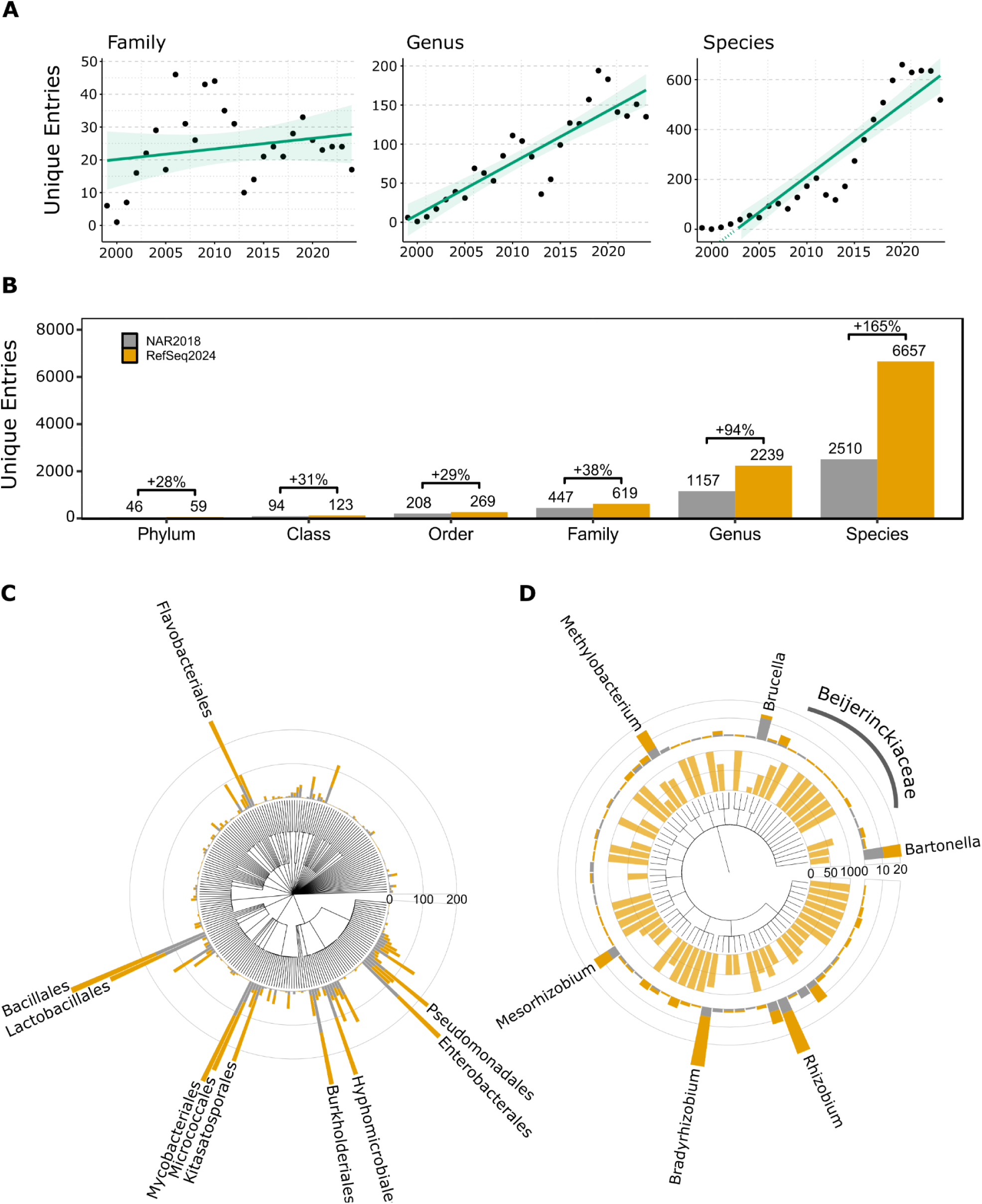
Increasing taxonomic coverage by complete RefSeq genomes. **A**. Number of distinct novel taxonomic ranks added per year, here shown for the taxonomic ranks family, genus and species, where a yearly increase can be noted. Data was fitted with a generalized linear model directly when plotting the data with the R package geom_smooth. **B**. A barplot with the respective count and percent increase of distinct entries for six taxonomic ranks from the NAR (Feb. 2018; gray) and RefSeq2024 (ochre/yellow) time stamps. Note: these numbers can slightly change over time, e.g. if taxonomic ranks get re-named or assemblies removed from RefSeq. **C**. A taxonomic tree visualized at the rank order with ggTree. The outer ring shows the number of distinct species for the NAR dataset (gray), and distinct new species added per order by RefSeq2024 in ochre/yellow. The top 10 orders with a large overall number of distinct species and/or a prominent increase in terms of number of distinct species added are labeled. **D**. Ggtree plot of Hyphomicrobiales, showing genera with many distinct species and/or a prominent increase in novel species added. The second ring shows percentage of newly added distinct species, the outer ring number of distinct species per genus.

Finally, we compared the RefSeq2024 dataset with an earlier reference time point from 2018(29), which we refer to here as the NAR2018 dataset (for short: NAR). A substantial increase in the coverage of different taxonomic ranks by complete genomes (**Figure 5B**) was noted: it ranged from 28% for the rank phylum (NAR: 46; RefSeq2024: 59) where 13 new phyla or candidate phyla were added, via 31% for class, 29% for order, 38% for family, 94% for genus to 165% for the rank species, where 4,147 new distinct species had been added (**Figure 5B**). We visualize this with the software ggtree (**Figure 5C**), capturing distinct taxonomic ranks recorded at the NAR time point (gray bars), here for the level species, and showing additional novel distinct species added by RefSeq2024 on top (ochre bars). For the rank order, the top ten orders in terms of novel distinct species added by RefSeq2024 are labeled (**Figure 5C**). They included Enterobacterales, which comprises many important pathogens, Pseudomonadales, that contain bacteria with important functions for plant growth promotion and biocontrol, Bacillales that include bacteria highly relevant for industry (enzyme production, fermentation) but also for agriculture (mainly from the genera *Bacillus*, *Paenibacillus* and *Brevibacillus*), including examples of biopesticides, i.e., *Bacillus thuringensis* and *Lysinibacillus sphaericus*), and finally Hyphomicrobiales (previously known as Rhizobiales), which include important soil microorganisms with roles as PGPB, as plant symbionts that fix atmospheric nitrogen and in biodegradation. A ggtree plot of the order Hyphomicrobiales visualizes the genera where most species have been newly added since the NAR dataset (**Figure 5D**): it implies the genera *Rhizobium* and *Bradyrhizobium* as hotspots, which is also visualized in a metacoder plot (**Figure S9B**). This plot, similar to the ggtree plot also allows to identify novel species with fewer assemblies that are linked to a higher taxonomic rank. Here, this is shown for example for the families Rhizobiaceae and Beijerinckiaceae, the latter of which includes free-living nitrogen fixing bacteria that do not rely on symbioses with plants to accomplish this, and methylotrophic genera that can use C1 compounds like methane and methanol which may help to reduce greenhouse gas emissions (**Figure 5D**). A metacoder plot for the order Lactobacillales, which includes diverse lactic acid bacteria that play important roles in the fermentation of various foods including cheese, is shown in **Supplementary Figure S11**. For the family Lactobacillaceae, we observed a substantial increase of well-studied genera (like *Lactobacillus*, *Limosilactobacillus* and *Ligilactobacillus*), but also many new genera for families where species were added that expand the taxonomic coverage of the GenomeCompendium, as for Carnobacteriaceae and Aerococcaceae.

### Exploring 16S rRNA gene copy number and sequence variation

The high-throughput analysis of variable regions of the 16S rRNA gene by amplicon sequencing has been widely used to describe the composition of natural microbial communities(62). The need to account for intragenomic variation of 16S rRNA gene copies has been raised(63), as it can lead to an overestimation of the operational taxonomic unit (OTU) count. Intragenomic 16S rRNA divergence was assessed in detail for 24,300 complete prokaryotic RefSeq genomes in 2023(64). We explored our roughly twice as big dataset using enrichment analyses, provide pairwise sequence identities for all 16S rRNA intragenomic gene copies that differ and explore other unique aspects, i.e., the preference of 16S rRNA genes for certain repeat types and quantifying how much of the repeat content of a genome is contributed by multiple copies of the entire rDNA operon.

We extracted 255,203 16S rRNA sequences from 46,720 RefSeq genomes (see Methods), the vast majority of which (99.8%) fell into a length range from 1450 −1600 nucleotides (nt) with a mean of 1539 +/− 19 nt. Overall, the 16S rRNA gene copy number ranged from 1 to 5 in archaea (data not shown) and 1 to 37 in bacteria (**Figure 6A**). A large drop in the total number per bin was observed for genomes with 9 or more copies, while genomes with 7 copies were most frequent. The assemblies with up to eight copies were enriched for the prominent genera *Escherichia*, *Klebsiella*, *Salmonella, Serratia and Shigella* (all from the order Enterobacterales), and assemblies from genera like *Pseudomonas*, *Streptomyces* and *Acinetobacter* (**Table S2**).

**Figure 6.**
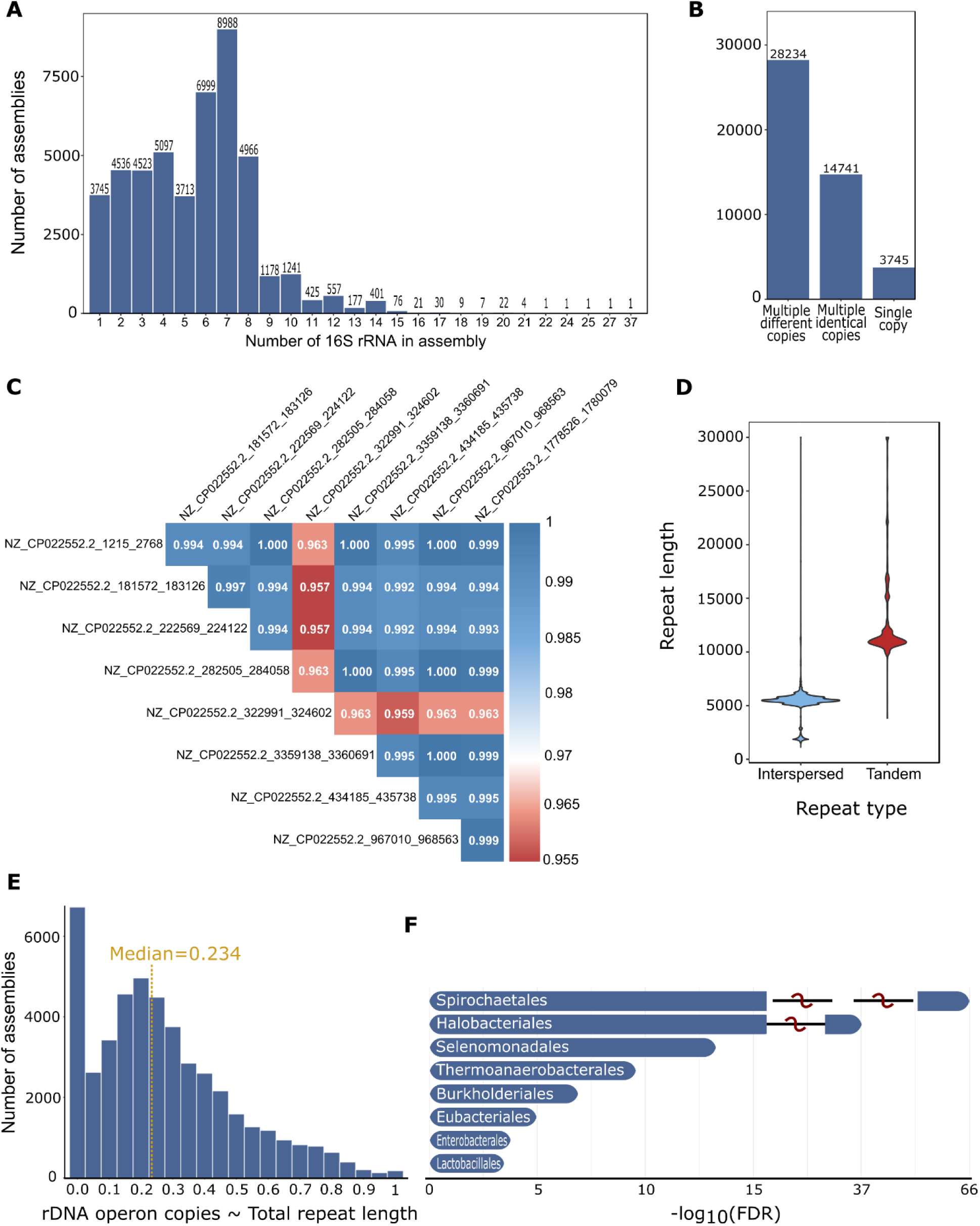
Analysis of 16S rRNA gene copy number, sequence variation and preference for repeat types. **A.** Histogram of the copy number of 16S rRNA genes in RefSeq genomes. **B.** The large majority of genomes have multiple 16S rRNA copies (92%), many of which can slightly differ in sequence. **C.** Pairwise sequence identity analysis among the nine intragenomic 16S rRNA gene copies of *Vibrio parahaemolyticus MAVP-R* (respective self-alignments are not shown). **D.** Length distribution of repeats containing 16S rRNA gene copies. Over 97% of 16S rRNA genes occur in interspersed repeats (blue), the remaining ones in tandem repeats (red), while none were detected in TIRs. **E.** Distribution of the ratio of the repeat region content contributed by rDNA operon copies over all RefSeq genomes with the median value marked (no partial overlap between repeated regions and operon coordinates were considered). **F.** Enrichment analysis at the taxonomic rank order for genomes that contain several rRNA genes, and where at least one of the 16S rRNA copies substantially diverges (similarity score < 0.97), evaluated using Fisher’s exact test and adjusted for multiple testing with the Benjamini–Hochberg method.

In contrast, among 3,745 assemblies with a single 16S rRNA copy (8.0%, **Figure 6B**), the genus *Mycobacterium* was significantly enriched (600 of 636 assemblies), which includes important animal (*M. bovis*) and human pathogens like *M. tuberculosis*, but also *Wolbachia* (292/293), *Rickettsia* (all 113) and *Borrelia* (all 78) (**Table S2**), i.e., three genera that represent endosymbionts or bacteria with a predominantly intracellular lifestyle, which are transmitted by arthropods and that feature small genomes and a reduced metabolism. The genus *Bradyrhizobium* with its nitrogen-fixing PGPB was also highly enriched (166/209 assemblies). Among RefSeq assemblies with 9 or more copies, the genera *Bacillus*, *Vibrio*, *Aeromonas* and *Paenibacillus* were highly enriched (**Table S2**), which correlates with the versatile habitats and environments that species of these genera (often motile) can live in, ranging from soil, water and aquatic habitats to animal hosts(65). Species of these genera also exhibit metabolic versatility (for some, this can include aerobic and anaerobic metabolism) and are able to thrive in different conditions, utilizing a variety of carbon sources.

We next explored the taxonomic enrichment for two important subsets, i.e., assemblies with multiple 16S rRNA genes with identical sequences (14,741 cases, 31.6%) and 28,234 assemblies (60.5%) with multiple copies, where for at least one of the pairwise alignments a sequence identity below 100% was observed (**Figure 6B**). Among the first subset, we found several genera for which over 90% of the assemblies had identical 16S rRNA sequences, including *Agrobacterium* (109/119), *Akkermansi* (100/110), *Xyllela* (76/83), *Phaeobacter* (60/62) and *Synechocystis* (14/14), (**Supplementary Table S2**). In contrast, among the genera for which most isolates had 16S rRNA copy sequence differences, we found *Vibrio* (637/660), *Paenibacillus*, *Fusobacterium*, *Dickeya* and *Morganella* (**Table S2).** Yet, most of these sequence differences were relatively small (greater 99% sequence identity). We thus also grouped the assemblies for which at least one copy had a sequence identity below 99% (3278 assemblies, **Table S2**) and below 97% (530 assemblies; **TableS2**). We plot the pairwise intra-genomic sequence identity, here for *Vibrio parahaemolyticus MAVP-R*which harbors 9 copies (**Figure 6C**), and provide similar figures in downloadable genome reports for RefSeq strains.

The large majority of 16S rRNA genes present in more than one copy in a genome were found in interspersed repeats; among 219,120 16S rRNA genes fully contained in exactly one repeat, 213,178 (97.3%) belong to this class (**Figure 6D**). Conversely, for 16S genes found in tandem repeats, we observed a signal of about twice the rRNA operon length (**Figure 6D**). We also computed how much the rDNA operons contributed to the overall repeat region content over all genomes: the median value of 23.4% indicates that repeat regions contain more genomic information than just rDNA operons (**Figure 6E**). Moreover, several orders including Spirochaetales, Halobacteriales and Burkholderiales were enriched for genomes where at least one of the intragenomic 16S rRNA copies had a sequence identity below 97% (**Figure 6F**). For ∼300 assemblies, the metadata analysis revealed that annotated rRNA genes were also localized on plasmids. We confirmed this for selected assemblies from the genera *Oecophyllibacter* and *Aureimonas* as reported before(66), and observed it for a substantial percentage of assemblies from *Priestia* (37 of 65) and *Azospirillium* (12/17). Finally, we found several cases where the annotations of chromosome and plasmid had been mixed up, i.e., another example where data mining efforts can help to improve quality control.

### Analysis of biosynthetic gene cluster content

Next, we explored preliminary identifications of potential biosynthetic gene clusters (BGCs) that can encode secondary metabolites with the protoclusters predicted by antiSMASH (see Methods) (**Supplementary Figure S12**). For simplicity, we here refer to the protoclusters as BGCs. They ranged from 0 to 98 hits per genome, with an overall mean of 9.5 and a median of 5 BGCs. Archaea harbored substantially fewer BGCs than bacteria, with a maximum of 13 BGCs (**Figure 7A**), but virtually the same average and median values as observed for bacteria. We then explored the distribution of BGCs for genera that play important roles for example in plant growth promotion and biocontrol (*Pseudomonas*) and as source of novel antibiotics (*Streptomyces*)(67). The genomes of both genera contained significantly more BGCs: *Streptomyces* with a mean of 48.6 (median 48) BGCs (in 1,253 RefSeq assemblies, **Figure 7B**), *Pseudomonas* with a mean of 19.3 BGCs (median 21) (2,031 assemblies, **Figure 7D**). Several genera exhibited a mean of ten or more encoded BGCs (**Supplementary TableS2**). These included *Kitasatospora* with 61, *Amycolatopsis* with 54, *Myxococcus* with 45, and *Saccharopolyspora* with 53 BGCs. Genomes of other genera with many assemblies but substantially lower median numbers of BGCs included disease related ones such as *Staphylococcus* with a median of 8 (2,085 assemblies, **Figure 7C**) or *Klebsiella* (9), *Escherichia* (6) or *Salmonella* with 4 BGCs, respectively. Besides observing relationships between the distributions of specific BGC types and certain genera, a classification analysis on a subset of the data revealed that the total number of BGCs per genome is consistently among the most informative features for predicting a genus, exhibiting a strong overall contribution to taxonomic classification alongside the presence or absence of individual BGC classes (**Figure 7E** and **Supplementary Results**). In addition, using the Apriori algorithm, we uncovered statistically significant co-occurrence patterns among BGCs across the analyzed genomes by applying association rule mining to tokenized sequences that preserved the binary presence or absence of individual BGC classes. Many of these rules were genus-specific, revealing conserved and characteristic BGC combinations; for example, tightly linked opine-like metallophore, hydrogen cyanide, phenazine, and naggn clusters in *Pseudomonas* (**Supplementary Figure S13**), or the predictive co-occurrence of a ni-siderophore cluster by cyclic-lactone-autoinducer and opine-like metallophore BGCs in *Staphylococcus* (for details, see **Supplementary Results** and **Supplementary Table S4**).

**Figure 7.**
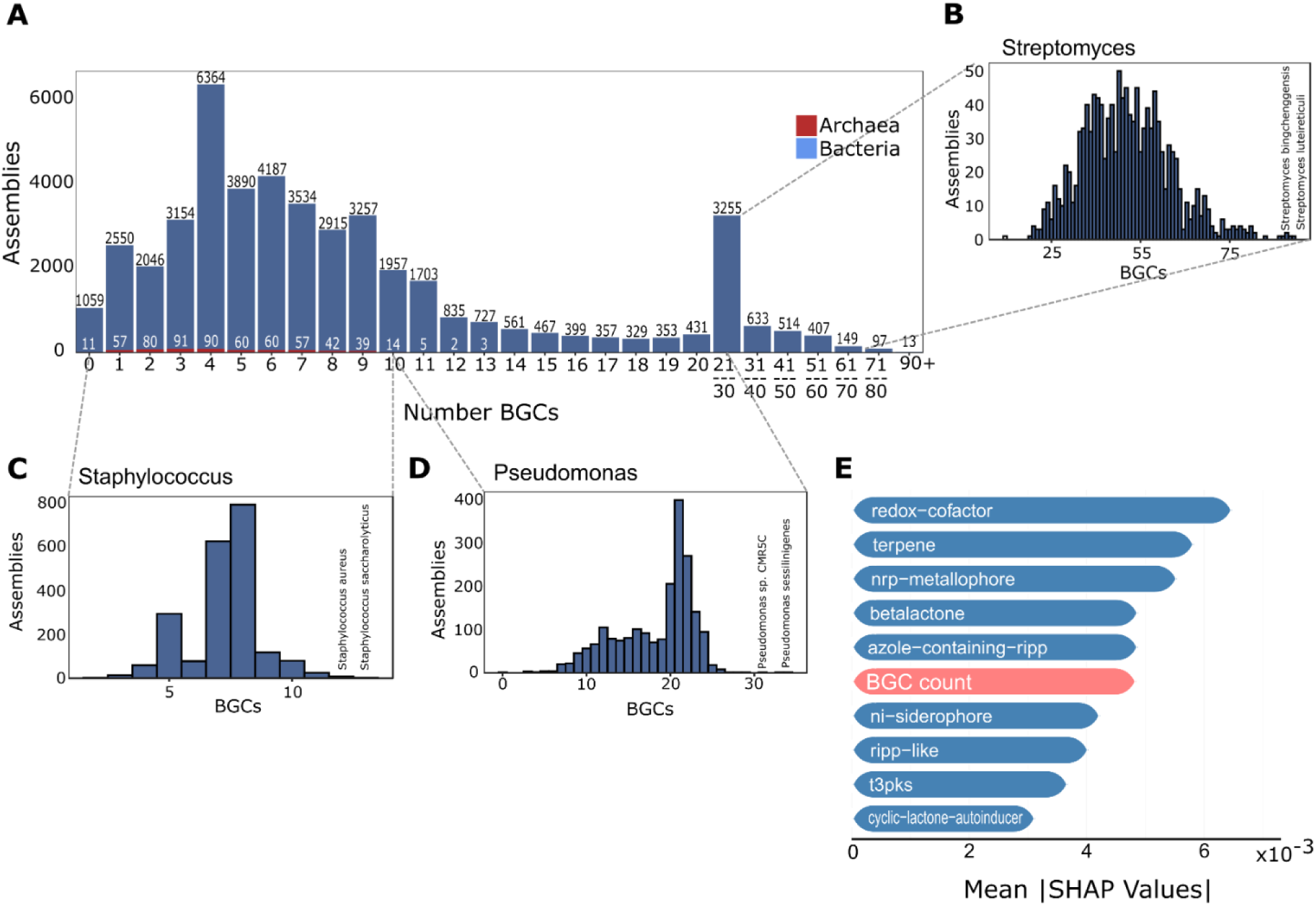
Analysis of biosynthetic gene cluster content. **A.** Histogram with the number of complete genomes (y-axis) per bin with the respective number of predicted BGCs (x-axis). For genomes with more than 20 BGCs, we used bins to group them. The BGC distribution for the genera *Streptomyces* (**B**), *Staphylococcus* (**C**) and *Pseudomonas* (**D**) is shown below. **E**. SHAP-based feature importance from a random forest classifier predicting genus-level taxonomy using 45 BGC classes and total BGC count. The total number of protoclusters per genome ranks among the top ten most informative features.

#### Selected use cases of the GenomeCompendium

By releasing the GenomeCompendium web server (https://genome-compendium.com/), we enable researchers to carry out several types of use cases (**Figure 8**): they can construct queries for interactive data mining (e.g. for quality control, genome assembly complexity, targeted sequencing into taxonomic ranks with low percentage of complete genomes, etc.), download detailed genome reports (**Figure 8A**), or execute a set of analyses on any user-provided complete prokaryotic genome using the GENARA web server (**Figure 8B**). By releasing ∼47,000 integrated proteogenomics databases (iPtgxDBs), we also support researchers to improve the genome annotation of their organism(s) of interest by proteogenomics (**Figure 8C**), i.e., using mass spectrometry-based proteomics data to identify missed CDS, a research field that has been gaining a lot of momentum (23, 25, 68).

**Figure 8.**
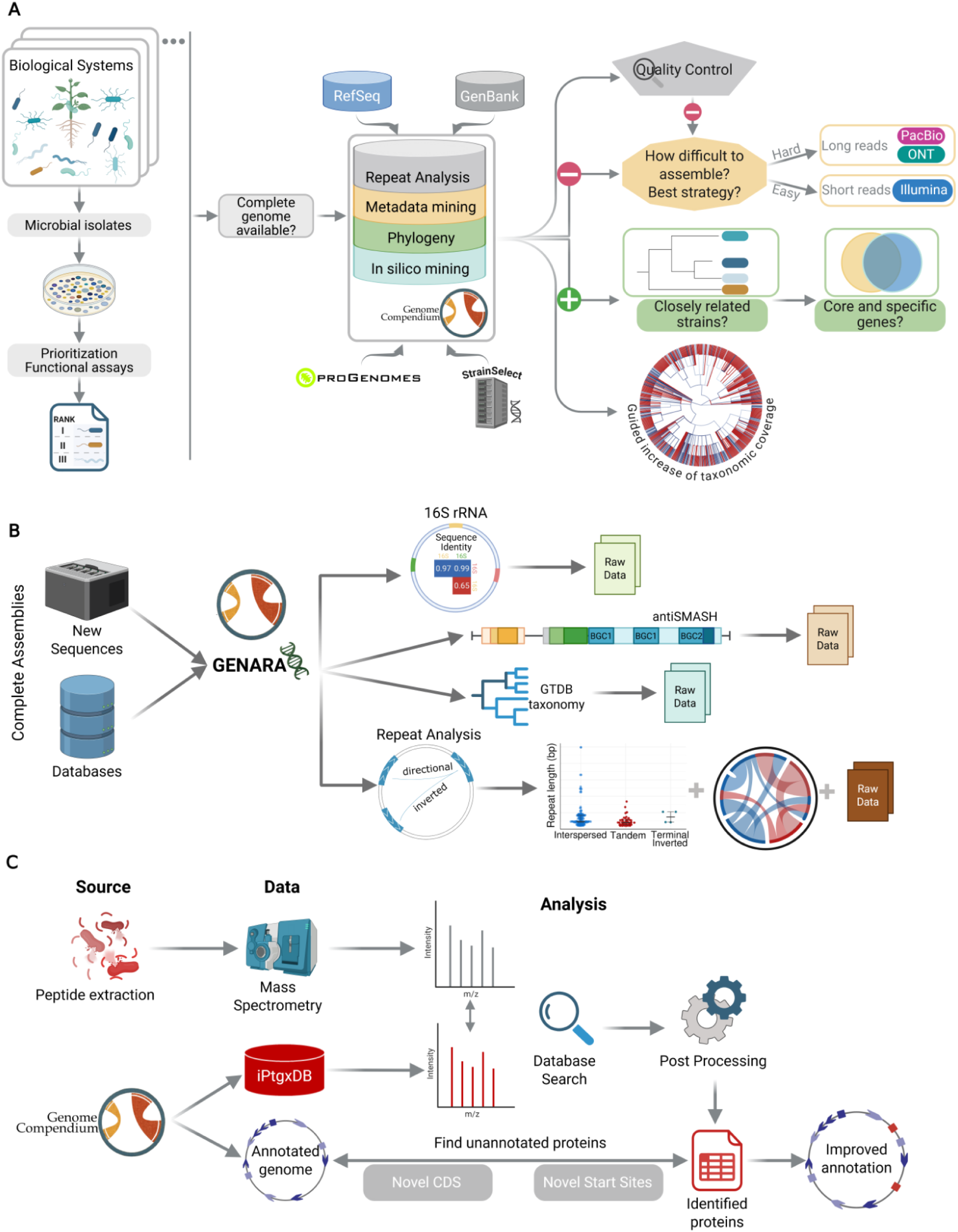
Selected use cases of the GenomeCompendium. **A**. Overview of selected data inquiries against the GenomeCompendium, typically starting with several prioritized microbiome isolates from any biological system of interest (see Results for more detail). **B**. Overview of the types of computations that can be run on any user-provided full prokaryotic genome sequence with the GENARA analysis tool. **C**. As a second unique feature, we provide close to 47’000 integrated proteogenomics databases for completely sequenced RefSeq strains, allowing researchers to find evidence for missed SEPs (red squares) and improve the genome annotation of their favorite model organism(s) using mass spectrometry-based proteomics data.

In the past, we regularly queried an in-house prototype of the GenomeCompendium to assess for which microbiome isolates of our experimental collaborators high quality, complete genomes had not yet been deposited at RefSeq. Ideally, these strains had been prioritized with functional assays, allowing us to sequence and de novo assemble the most relevant isolates(69) (**Figure 8A**). Prioritizing strains in this manner will increase the taxonomic coverage and novelty. Conversely, for strains where a complete genome exists, users can now check whether our QC steps have flagged it, i.e., indicating a reference-based assembly strategy has been used or the assembly contains long repeats but the metadata indicates it has been assembled using only short read data (**Figure 8A**, right panel). In both cases, users can opt to generate a new, high quality assembly using long reads (and short reads to polish). Moreover, the assembly complexity of available strains of a taxonomic rank (e.g. species or higher) can be checked to be able to adapt the sequencing strategy, for example to include a size selection step if a high percentage of complex class III genomes exists, or to additionally rely on very long reads from the ONT platform, as we had done to resolve a particular complex genome(29). Importantly, for isolates where other closely related, completely sequenced strains exist, phylogenetic trees can be constructed and comparative genomics approaches carried out on phylogenomic subsets to identify core, accessory and strain specific genes (**Figure 8A**) and to use this as a basis to link phenotypic observations to genotypic differences; for an example of this use case, see (69).

On the website, a zoomable Figure shows genera for which no or only few complete genomes exist, supporting targeted sequencing efforts (**Figure 8A**). Notably, large-scale data downloads are supported via programmatic access: a dedicated download handler allows users to retrieve data for any assembly directly using its accession, while the full data table can be explored and mined interactively through the web interface. To further enhance its value, we provide crosslinks for the complete genomes to the proGenomes resource(19) where available, allowing researchers to leverage their accurate and consistent genome annotations(70) (see **Table S1**, **Figure S2**). We also link a subset of Refseq and GenBank assemblies with StrainSelect identifiers (**Supplementary Figure S14 and Table S1**), which provide information where a strain can be purchased from(45).

Researchers can analyze their own completely sequenced genomes to extract results from four types of analyses through the GENARA tool (**Figure 8B**). They can i) identify repeats in their sequences (using the same parameters as in our analyses) and visualize them with summary plots, enabling detection of potential quality issues and genome assembly difficulty; ii) perform 16S rRNA identification when annotations are missing, providing sequence identity scores for multiple intragenomic 16S rRNA copies, which is helpful to estimate the accuracy of amplicon sequencing based OTU counts; iii) assign GTDB taxonomy, allowing standardized classification and comparison across genomes and with respect to reference datasets; and iv) access raw antiSMASH output files, facilitating exploration of BGC diversity, co-occurrence analysis, functional potential, and specialized metabolite predictions.

Finally, as another unique aspect of the GenomeCompendium, we democratize the ability of researchers to improve the genome annotation of their favorite prokaryote(s). This is highly relevant: in *Escherichia coli*, arguably the best studied model bacterium, over a span of ten years researchers identified and experimentally validated ∼140 CDS encoding proteins smaller than 50 aa(71). To find evidence for often missed short ORF encoded proteins (SEPs) that can carry out important functions, we release a custom, small iPtgxDB(23) (47) for each of the 46,754 RefSeq assemblies (downloadable from https://genome-compendium.com/). The iPtgxDBs consolidate multiple annotation sources (RefSeq, an ab initio prediction (Prodigal) and candidates from GMSC-mapper(50); see **Supplementary Figure S15**) into a minimally redundant search database. Thereby, almost all peptides identified in proteomic data can be unambiguously matched to unique protein entries, simplifying downstream analysis and enabling the discovery of novel CDS and alternative start sites. The identification of such SEPs has become the focus of intense research efforts (reviewed in(25, 72, 73)).

## Discussion

The exponential increase of prokaryotic genome assemblies, both for bacteria, and -at a much lower absolute rate-for archaea (**Figure S1A**) has been fueled by advances in sequencing technologies and assembly algorithms and by declining sequencing costs (74). While the vast majority of genome assemblies are still based on short read data, availability of long-read data significantly improved the number of complete prokaryotic genomes (**Figure 1A**, **Figure S1C**). As previously shown, complete genomes represent an optimal basis for downstream functional genomics, comparative genomics and systems biology studies(26, 75), e.g. to arrive at a mechanistic understanding of an antagonistic activity against a pathogen or the mode of action of a new antibiotic(76). Completely sequenced strains that can be procured allow researchers to create, test and iteratively improve SynComs that combine beneficial aspects of several strains and to translate insights from microbiome research into practical applications in agriculture(18), biotechnology(77) or medicine(78). We use the complete genomes as a basis to integrate and leverage available metadata and in-house computations (**Figure S2**), such as the repeat analysis, the intragenomic 16S rRNA variation and proteogenomics databases for almost 47,000 complete RefSeq genomes.

By focusing on complete genomes, we miss out on the value of metagenomic field studies and metagenome assembled genomes (MAGs), which add genome sequences of novel species and functional insights into as of yet unculturable prokaryotes(13, 14). Sampling of terrestrial geothermal springs, for example, substantially expanded the known phylogenetic diversity of archaea, adding close to 400 newly identified species from nearly 3,000 reconstructed MAGs(79). However, a drawback of MAGs is that researchers do not have an isolate in hand that they can study under controlled experimental conditions. Moreover, MAGs are already comprehensively represented in valuable genomic resources such as the GTDB(80).

The presence of repeats is the main obstacle for complete genome assembly. We focus on the frequent longer and near-identical repeats (≥ 500 bp with sequence identity ≥ 95%) that complicate *de novo* assembly of prokaryotic genomes(28), but acknowledge that shorter repeats can also be highly relevant. In methane-oxidizing archaea for example, large Borg elements (linear extrachromosomal elements flanked by TIRs) contain tandem direct repeats in ORFs which are under strong selective pressure and often found in intrinsically disordered regions of membrane and extracellular proteins(81). In *Mycobacterium tuberculosis* strains, the GC-rich PE/PPE (proline, glutamic acid; proline-proline-glutamic acid) family members harbor short repetitive sequences that have long represented genomic blind spots despite the relevance of this gene family for virulence, host pathogen interaction and thus as candidates for the development of novel diagnostics, vaccines or therapeutics(82).

By integrating metadata with our repeat analyses, we uncover that up to 6.3% of the analysed genomes may harbor assembly errors or miss sequence information (**Figure 4**); these QC aspects can be readily identified through interactive data mining through the web interface. Moreover, we provide proof that over time, increasingly complex prokaryotic genomes could be resolved (**Figure 3A**), for which long read sequencing data was critical (**Figure 1C**, **Figure S1B**). The overall complexity of prokaryotic genomes has thus been substantially underestimated (**Figure 3**, **Figure S7**), as evidenced by the much higher percentage of complex class III genomes with repeats over 30kb, i.e., 5,8% compared to 3.2 % observed until 2018(29). Integration of taxonomy classifications from NCBI and GTDB with the genome assembly complexity metric allow us to flag taxonomic ranks that are particularly difficult to assemble. This creates a basis for targeted sequencing aimed at increasing the taxonomic coverage of completely sequenced genomes (**Figure 1E**, **Figure 3C, D**). Comparing different snapshots over time allows researchers to identify novel taxonomic ranks that capture interesting biology, including PGPBs relevant for sustainable agriculture, lactic acid bacteria for fermented foods and methylotrophic soil bacteria that may help to reduce climate gas emissions (**Figure 5, Figures S9, S11**) to name a few.

Additional biologically relevant aspects include the analysis of i) AMR gene classes within repeat regions for selected genera (**Figure 2**) including the ESKAPE pathogens **(Figure S6**), ii) the copy number and sequence variation of multiple intragenomic 16S rRNA gene copies (**Figure 6**), which forms the basis to link amplicon sequencing variants (ASV) data to complete genome information and correct inflated OTU counts (63), and of iii) BGCs (**Figure 7**), which capture the potential of strains to produce secondary metabolites and many other relevant products. Genomes that feature high BGC counts often serve as a source for the identification of novel antibiotics, as has been reported for the genus *Kutzneria* (median BGCs: 79)(83). The addition of 782 *Streptomyces* strains in 2024 -readily identifiable by analysis of the integrated metadata-likely also reflects the push to identify strains that produce urgently needed novel antibiotics(84) - about two thirds of clinical drugs were isolated from *Streptomyces* species(67) - but also to utilize their potential for applications in agriculture(85, 86). Analyzing the enrichment of such features for specific taxonomic ranks -one important use case (**Figure 8**) - can uncover interesting biology such as an enrichment of transposases that can lead to reductive genome evolution (87). This became evident, for example, by the significant enrichment of class II genomes at the species level among *Bordetella pertussis* (595 of 633 strains) but not *B. parapertussis* (0 of 92 strains), with the former amassing a median of 259 IS481 insertion sequence copies, validating previous findings (88). The ability to place completely sequenced genomes into the phylogenomic context and leverage comparative genomics approaches, ultimately linking phenotype to underlying genotypes is highly relevant, as is the addition of new species with complete genomes guided by targeted sequencing (**Figures 3, 8**), which can increase their taxonomic coverage and add value for studies that rely on SynComs.

The GenomeCompendium represents a reliable reference for the growing set of complete prokaryotic genomes for comparative analyses, in silico data mining and various other use cases. As such, it provides the infrastructure necessary for data integration, reproducibility, and the systematic exploration of biological systems, here of prokaryotes. All data, including cross-references to valuable other resources like GTDB, proGenomes and StrainSelect is available to users, either by downloading extensive reports for their genome(s) of interest or interactively mining the GenomeCompendium web server. As two unique features, complete prokaryotic genomes can be uploaded and analyzed with the GENARA analysis tool, and precomputed iPtgxDBs for any of the ∼47k RefSeq genomes can be used to improve their genome annotations with proteomics (24, 82), and serve as a basis to dive into unannotated small proteins, which represent the other end of the spectrum of the dark matter of prokaryotic genomes (10, 89). Beyond this, the GenomeCompendium constitutes a well-structured substrate for data mining(90), supporting functional genomics and discovery-driven studies, including approaches that couple large-scale genome mining with conversational interfaces in which AI models resolve free-text requests into structured queries over the ∼90 annotated features, allowing users in the future to identify the best-suited genomes for various applications.

## Acknowledgements

We thank Dr. Philipp Hugenholtz and his team for access to the GTDB, Dr. Daniel Mende and his team for help with cross-referencing to proGenomes and Dr. Todd deSantis for providing access to the StrainSelect dataset. Further, we thank Joelle Sasse (Agroscope), Gabriella Pessi (University of Zurich) and Wolfgang Hess (University of Freiburg) for helpful suggestions and feedback on the manuscript, and German Bonilla-Rosso (Agroscope) for helpful suggestions on the web server functionality. Parts of the figures were designed in BioRender.

## Data Availability

Raw data, as well as compiled and raw analysis results are freely available without registration at https://genome-compendium.com/. The analysis pipelines comprising GENARA are available on ZENODO(30) and at https://github.com/mgbteam/GENARA. The GenomeCompendium and GENARA are released under a GPLv3 licence.

