## Supplemental Information for "GenomeCompendium: A database for the integrated analysis of repeats, assembly quality and functional content of complete prokaryotic genomes"

### Supplementary Information

#### Index

#### Supplementary Methods

Repeat analysis - We detect and classify four types of repeat sequences in prokaryotic genomes using the public tools Nucmer v 3.1(1) and equicktandem, before integrating and further simplifying the output.

Interspersed repeat analysis - To identify interspersed repeats, Nucmer was run (parameters –maxmatch –nosimplify (self-alignment)) and results filtered (command delta-filter –l500 –i95) to retain repeats ≥ 500 bp and with ≥ 95% sequence identity; self-alignments on the main diagonal were discarded (**Supplementary Figure S3**, Step 1). The directionality of the repeats on the same replicon was added as '*inv*', '*dir*' (inverted repeat or directional repeat) and as 'inter' for repeats located on different chromosomes or plasmids. The list of repeat pairs was further simplified by custom scripts in 3 steps (**Supplementary Figure S3**, Step 2). Next, we split repeat pairs into single repeats and combined them into so-called repeat regions: repeats which overlap by more than 95% of the smaller repeat as well as repeats that are completely contained within another repeat are fused to one repeat region (Step 3). The output of this step is compared to other repeat types as described below and in **Figure S3** (Step 6). If an interspersed repeat region is completely contained within a tandem or a terminal inverted repeat (see below), it was deleted from the list.

Tandem repeat analysis - Two strategies were used to identify tandem repeats: First, we ran the software equicktandem with –maxrepeat 10,000 and –threshold 100 and filtered the output for tandem regions ≥ 500bp and with ≥ 95% sequence identity. In a second step, the repeat pairs identified in the interspersed repeat analysis are categorized as tandem repeats, provided they are direct repeats (positioned within 10 bp of each other) or directional repeats with an overlap of 95% or less (see **Figure S3**, Step 4). The output is also compared to the list of interspersed repeat regions (see **Figure S3**, Step 6). All tandem repeats which are completely contained within a larger interspersed repeat are deleted.

Terminal inverted/terminal directional repeat analysis - For all linear contigs, we carried out an additional terminal inverted repeat search (see **Figure S3**, Step 5): Repeat pairs where both repeats are localized at the end of a linear contig (with a flexible start of up to 20 bp from a terminus) and are inverted are classified as terminal inverted repeats (TIR). Directional repeat pairs at the end of linear contigs (with a flexible start of up to 20 bp from a terminus) are classified as terminal directional repeats. They are treated as a special case as some of them may indicate the lack of a circularisation step for a chromosome or plasmid. Finally, the output files of individual repeat analyses were merged into a single file (<assembly>_ALL_REPEATS.gff).

AMR gene enrichment analysis in repeats - The Metafor R package(2) was used to perform a Mantel-Haenszel (MH) enrichment using species as strata and the log odds ratio as a measure. The odds ratio is the ratio of how often an analyzed feature was found within identified repeats (requiring at least 70% overlap with a repeat) compared to all other background genes. Due to the bias of overrepresented species, a random-effects (RE) estimate was computed as well on the per-stratum log odds ratios, on which a continuity correction of 0.5 was applied on strata with at least one zero cell. The between-species variance was estimated through restricted maximum likelihood, and the confidence intervals and tests used the Knapp-Hartung adjustment. When the restricted maximum likelihood failed to converge, the Paule-Mandel estimator was used. Heterogeneity was quantified by the estimated between-species standard deviation of log odds ratios (tau), the $I^2$ statistic, and Cochran's Q p-value. A feature was considered significant at the aggregated or individual genus level if the absolute $log_2OR$ was higher than 2 and both Benjamini-Hochberg-corrected MH p-values and RE p-values were below 0.05.

Proportion of complete genomes - Plots showing the proportion of complete genomes for a given taxonomic rank were generated using a number of additional R packages including *data.tree* (3), *ape* (4), *ggtree* (4, 5), *igraph* (4–6), *viridis* (7), *MicrobiotaProcess* (8), *ggtreeExtra* (9) and *treemapify* (9, 10). These packages were used for data manipulation, visualization, tree construction, and hierarchical plotting to effectively represent taxonomic completeness (**Figure 1**) and assembly complexity across different ranks (**Figure 3**).

NCBI taxonomy information processing - We retrieved species information from the NCBI taxonomy using python's ete3 library v 3.1.3 (11), see column "Species(NCBI)" (**Table S1**). We added a column "Species_defined(NCBI)" (our edited data), for which we required a full scientific name with the two tags for genus and species. If the species name began with "Candidatus", the first three words (Candidatus genus species) were retained. Names containing the strings "archaeon", "bacterium", "sp.", "genomosp.", "genosp.", "phylum", "haloarcheon", "proteobacterium", "endosymbiont", "symbiont", or "phytoplasma" were manually assessed; they did not represent true species names (including a few cases where the species name of the host of a symbiont was given, or of a higher taxonomic rank) and thus not retained in our edited data. Overall, 7,526 assemblies (5,645 from RefSeq) did not have a "species_defined" name. For 26 RefSeq assemblies no genus level info was provided; they were not considered for selected enrichment analyses (see separate sheets, **Table S2**).

rRNA operon analysis - The coordinates of full rRNA operons were extracted using Barrnap (Basic ribosomal RNA predictor) version 0.9 (12) by identifying the genomic locations of the 5S, 16S, and 23S rRNA genes across all assemblies. Only RefSeq assemblies with a valid genus-level taxonomic identifier were retained for this analysis (46,728). For each genome, the set of identified rRNA gene coordinates was cross-referenced with annotated repeat regions. A repeat region was considered to contain a full rRNA operon if it overlapped all three rRNA genes - 5S, 16S, and 23S - based on coordinate comparison. These overlaps were used to flag repeat elements that encompassed complete rRNA operons. For each genome, the cumulative length of repeat regions containing a full rRNA operon was computed and expressed as a proportion of the total repeat length.

16S rRNA divergence analysis - Pairwise 16S rRNA gene similarity values from intragenomic 16S rRNA copies were imported from VSEARCH (13) output files. Assemblies with any pairwise identity below 97% were flagged as divergent, and the proportion of divergent comparisons per assembly was calculated. Taxonomic metadata from phylum to species levels was merged with the divergence data. Enrichment of divergent assemblies within each taxonomic rank (phylum, class, order, family, genus, species) was assessed using Fisher's exact test. For each taxonomic group, the number of divergent and non-divergent assemblies within the group was compared to those outside the group. P-values were adjusted using the Benjamini-Hochberg method to control for multiple testing. The analysis was performed only for RefSeq assemblies and for assemblies with all the taxonomic ranks annotated.

BGCs: Genomic text mining, association rule mining and genus-level enrichment analysis through interpretable supervised learning - This analysis was performed on RefSeq assemblies with valid genus-level taxonomic assignments (46,728). Genome sequences were segmented into 100 bp blocks, each encoded using a two-character identifier representing a specific BGC product. These identifiers were concatenated into a collapsed, tokenized string per genome. Positional information and feature length were omitted to simplify the model and focus on the presence of specific BGC types, which is typically sufficient for classification and co-occurrence tasks. However, this decision may limit the ability to capture spatial relationships between co-localized BGCs, which could provide additional biological insights into their functional associations. Association rule mining was conducted using the Apriori algorithm implemented in the arules R package (14). Rules were identified using a minimum support threshold of 1%, a minimum confidence of 80%, and a rule length between 2 and 4 items. Supplementary Table S4 lists all rules with a lift greater than 2, considered indicative of non-random co-occurrence between features. Genus-specific rule enrichment was tested using a Fisher's exact test with Benjamini-Hochberg correction for multiple comparisons. Only genera with at least 10 assemblies were considered, and associations with a false discovery rate (FDR) below 0.01 were deemed significant. To assess the predictive relevance of BGC-related features, a random forest classifier with 300 decision trees was trained on the tokenized genome sequences using a stratified five-fold cross-validation split, with each fold composed of 80% training and 20% testing data. Class labels corresponded to genus-level taxonomic assignments, and only genera with at least 100 entries were retained (66 genera) to ensure statistical robustness and faster computation. BGCs present in fewer than 1% of the assemblies were not considered. Tokenized genome features were treated as binary variables without additional transformation. The number of BGCs per assembly was added as an additional feature. Model performance was evaluated using mean classification accuracy and macro-averaged F1-score across folds. A multinomial logistic regression model was trained on the same data split and used as a naïve baseline for performance comparison. Feature importance was assessed using SHAP (SHapley Additive exPlanations) values (15) derived from the trained random forest model. SHAP values were aggregated as the mean absolute Phi value per feature across all test instances to provide a global ranking of the most informative genome tokens contributing to genus-level classification. The following R packages were used in the analysis: *arules* (14), *randomForest* (16), *car* (17), *data.table* (18), *dplyr* (19), *glmnet* (20), *doParallel* (21), *purrr* (22), *caret* (23), *iml* (24), *readr* (25), *stringr* (26), *ggplot2* (27), *tidyr* (28), *stats* (29) *and Cairo* (30).

GenomeCompendium public web server - The platform was developed as a Golem Rshiny package (31) and relies on several additional R packages. These included shiny (32), shinycssloaders (33), plotly (34), shinydashboard (35), shinyWidgets (36), bslib (37), shinyBS (38), shinyjs (39), DBI (40), RPostgres (41), pool (42), promises (43), future (44), DT (45), callr (46), uuid (47), and stringr (26).

#### Supplementary Results

##### Summary over longest repeats and range of occurrences per repeat type.

The longest repeat pairs among 46,092 genomes with 1-1,323 interspersed repeats could span up to 1584 kb, among 42,074 genomes with 1-163 tandem repeats they could span up to 1426 kb (**Figure S5A**). The distribution of the maximum repeat length for 41,853 genomes with both interspersed and tandem repeats is shown in **Supplementary Figure S5B**. In contrast, terminal inverted repeats (TIRs) spanned up to 2415 kb but were only found in 741 genomes (overall 1,690 repeats, 0.1%) ranging from 2-10 copies; their length distribution is shown in **Supplementary Figure S5C**. Terminal directional repeats were smaller (up to 87.8 kb) and found in 191 genomes with 2-12 copies (overall 470).

##### Co-occurrence analysis of biosynthetic gene clusters

To assess whether statistically significant co-occurrence patterns exist among BGCs products identified in the analyzed genomes, an association rule mining approach was applied to tokenized genome sequences that preserved the binary presence or absence of individual BGC classes (see Methods). The Apriori algorithm was used to identify co-occurrence relationships among BGC types. The resulting frequency distribution (**Supplementary Figure S12**) revealed a high prevalence of nonribosomal peptide (nrp) and ribosomally synthesized and post-translationally modified peptide (RiPP) BGCs, along with terpene-precursor clusters. Using stringent criteria (lift > 10, confidence > 0.95, and support > 1%), several robust association rules emerged (**Supplementary Figure S13**). Notably, strong dependencies were identified among opine-like metallophore, hydrogen cyanide, phenazine, and naggn BGCs. Enrichment analysis at the genus level demonstrated that these associations were highly specific and strongly enriched in Pseudomonas (**Supplementary Table S4**). Bacillus also exhibited distinct and characteristic co-occurrence patterns, including frequent rules linking betalactone, nrps, sactipeptide, and type III polyketide synthase BGCs. For instance, in 491 of the 1,793 Bacillus assemblies analyzed, the concurrent presence of nrps, t3pks, and sactipeptide BGCs reliably predicted the additional presence of a betalactone cluster; this combination that occurred in only four non-Bacillus assemblies. Additional genera including Streptomyces, Staphylococcus, Acinetobacter, and Mycobacterium were similarly defined by highly specific BGC co-occurrence rules. While some of these genera are known to encode a high median number of BGCs per genome, facilitating the emergence of such patterns, Staphylococcus and Acinetobacter are particular exceptions, with average BGC counts of seven and nine, respectively. Staphylococcus genomes displayed a particularly prominent rule as in 1,870 of 2,085 assemblies, the co-occurrence of cyclic-lactone-autoinducer and opine-like metallophore BGCs was predictive of a ni-siderophore cluster. This rule was otherwise observed in only 59 genomes outside Staphylococcus, being statistically significant only for Brevibacillus and Paenibacillus.Given that the most distinctive BGC classes are often observed in genomes with above-average BGC abundance, we investigated the extent to which total BGC count per assembly contributes to taxonomic classification at the genus level. To this end, we employed an interpretable supervised learning framework using a random forest classifier trained on the binary presence or absence of 45 BGC classes, supplemented by the total number of BGCs per genome as an additional quantitative feature (see Methods). The model was evaluated using five-fold cross-validation with an 80:20 training-to-testing split, yielding a mean classification accuracy of 0.85 and a macro-averaged

F1-score of 0.75 across the folds. These results markedly outperformed a baseline logistic regression model, which achieved an accuracy of 0.75 and a macro-averaged F1-score of 0.50, underscoring the added value of non-linear interactions in capturing the taxonomic signal. To gain insight into the relative importance of individual predictors, we applied SHAP (SHapley Additive exPlanations), a model-agnostic feature attribution method (see **Supplementary Methods**). Among the 46 predictors, the total number of BGCs consistently ranked among the top most informative features (**Figure 7E**), with SHAP values nearly equivalent to those of the highest-ranked BGC types. This result underscores the biological and taxonomic relevance of BGC abundance as a predictive and interpretable feature for genus-level classification.

#### Supplementary Figures and Tables

**Table S1.** Master table.

This “Master” table summarizes metadata extracted from summary files of NCBI RefSeq (46,754) and additional GenBank (13,224) assemblies, and that shows the results of several computations. Please see the legend sheet as it provides a detailed overview and description of the data in different columns of this extensive table, which we have often used for exploratory data analysis and mining.
**See separate Excel file**.

**Figure S1**. **Increase of complete genome assemblies at NCBI Refseq and GenBank.**

**A**. The cumulative increase of the number of complete genomes over time (2001-2024) is substantially slower for archaea (orange) compared to bacteria (light blue); for the data points from 2001-2007, 2007-2014 and 2014-2024 separate linear regressions on log-transformed data were carried out (fitted orange and blue lines, respectively) to capture significant decreases in sequencing costs (grey line). **B**. Barplot with the number of complete genomes added per year (not cumulative) and the respective long read sequencing technology used: PacBio (green), ONT (red). **C**. The relative percentage of complete genome sequences deposited in RefSeq has been decreasing each year until 2016 (8.3%). This can be attributed to the fact that more assemblies of lower completeness levels, i.e., scaffolds (light gray) and contigs (dark gray) were deposited. From 2017 on, this trend is slowly reversed as a higher percentage of complete genomes got deposited that rely on long read sequencing data (see **Figure 1D**), reaching 12.7% complete genomes in 2024 (the actual percentage may differ slightly, as no metadata was made available for 9.5% of the assemblies). **D**. The trend of the decreasing fraction of complete genomes deposited each year is more pronounced (and not reversed) for GenBank assemblies, as the vast majority got deposited in the form of contigs (10s-100s per assembly), particularly from 2019 onward (dark gray bars). By the end of 2024, contig level assemblies accounted for ~82.7% of the 2,453,279 GenBank assemblies, and only ~2.4% were complete genomes. **E**. Cumulative increase of the four assembly categories over time in GenBank. By the end of 2024, 59,978 complete genomes were available, i.e., 13,224 genomes more than from RefSeq. **F**. The Sankey plot illustrates the main categories of the 13,224 additional complete assemblies only deposited in GenBank (subgroups with at least 20 assemblies are shown). Note: a strain can belong to more than one category. Major fractions of these additional genomes stem from i) large multi-isolate sequencing projects of pathogens (over 7300, including the genera *Escherichia*, *Klebsiella, Helicobacter, Staphylococcus*, *Salmonella*, *Enterococcus*) that are accepted by GenBank but not by RefSeq, ii) most assemblies from metagenome projects (although RefSeq recently started to include selected cases that pass their quality control criteria), iii) genomes with many frameshifted proteins, iv) of undefined genus or unverified source origin, as well as v) contaminated samples to name the most prominent ones.

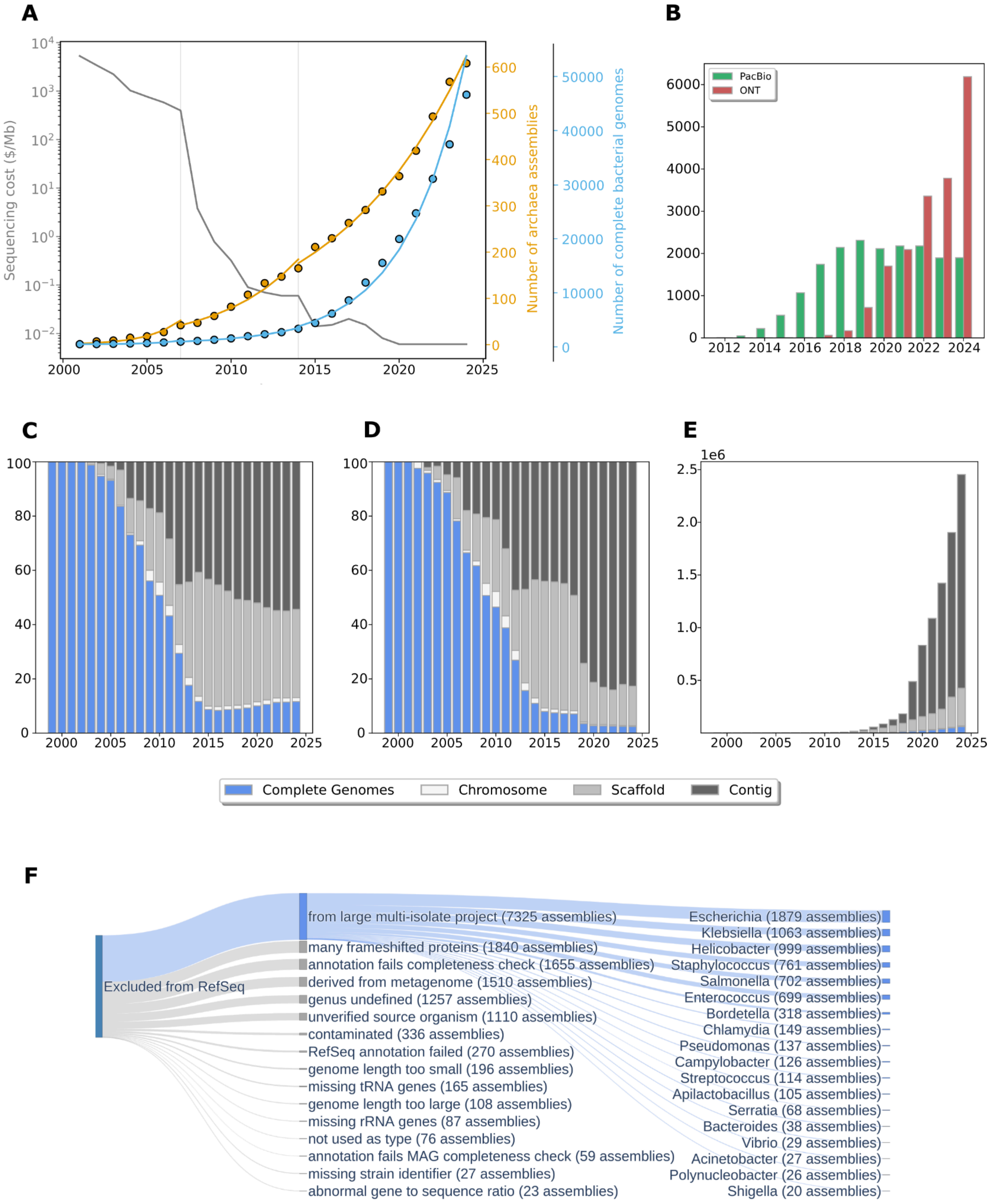

A
Sequencing cost ($/Mb)
Number of archaea assemblies
Number of complete bacterial genomes
B
PacBio
ONT
C
D
E
1e6
Complete Genomes
Chromosome
Scaffold
Contig
F
Excluded from RefSeq
from large multi-isolate project (7325 assemblies)
many frameshifted proteins (1840 assemblies)
annotation fails completeness check (1655 assemblies)
derived from metagenome (1510 assemblies)
genus undefined (1257 assemblies)
unverified source organism (1110 assemblies)
contaminated (336 assemblies)
RefSeq annotation failed (270 assemblies)
genome length too small (196 assemblies)
missing tRNA genes (165 assemblies)
genome length too large (108 assemblies)
missing rRNA genes (87 assemblies)
not used as type (76 assemblies)
annotation fails MAG completeness check (59 assemblies)
missing strain identifier (27 assemblies)
abnormal gene to sequence ratio (23 assemblies)
Escherichia (1879 assemblies)
Klebsiella (1063 assemblies)
Helicobacter (999 assemblies)
Staphylococcus (761 assemblies)
Salmonella (702 assemblies)
Enterococcus (699 assemblies)
Bordetella (318 assemblies)
Chlamydia (149 assemblies)
Pseudomonas (137 assemblies)
Campylobacter (126 assemblies)
Streptococcus (114 assemblies)
Apilactobacillus (105 assemblies)
Serratia (68 assemblies)
Bacteroides (38 assemblies)
Vibrio (29 assemblies)
Acinetobacter (27 assemblies)
Polynucleobacter (26 assemblies)
Shigella (20 assemblies)

**Figure S2. Overview of the metadata extraction, repeat detection, feature analysis and cross referencing workflow.**

The subset of complete prokaryotic genomes is downloaded from NCBI (RefSeq and GenBank; incremental updates). Metadata is extracted, augmented with the results from several computations and integrated in the summary table and individual report files. A repeat detection workflow is run on the extracted fasta files, which relies on Nucmer(1) and equicktandem (part of the EMBOSS open software suite); subsequent simplification steps lead to a compilation of lists of interspersed, tandem and terminal inverted repeats and repeat regions (for details, see **Figure S3**; their respective colors are also used in the report files, see below). The repeat lists are also made available as GFF files to be loaded into a genome browser. Additional feature analyses are carried out, like an analysis of the copy number and sequence divergence of 16S rRNA genes, and an antiSMASH analysis for the prediction of biosynthetic gene clusters (BGCs)(48). The results of the GTDB taxonomy(49) are extracted (or computed) and compared to that from the NCBI and stored in the Summary table. Crosslinks to valuable resources like proGenomes(50) with its more detailed annotation and StrainSelect(51), which lists vendors where selected strains can be purchased from, are made available for those entries where such a cross-link exists (22,862 of the complete RefSeq and 4,235 of the complete GenBank genomes could be linked to a proGenomes entry. 4,071 of 46,754 Refseq assemblies and 199 of the 13,224 additional GenBank assemblies could be linked to a bioresource listed by StrainSelect). This wealth of data is integrated in a Summary table ("Master" **Table S1**) and in individual report files for each of the analyzed assemblies (PDF format). These data resources represent a valuable basis for data mining and knowledge discovery. To democratize the ability of researchers to further improve the genome annotation of their favorite prokaryote(s), in particular to find evidence for often missed short ORF encoded proteins (SEPs) that can carry out important functions, we release a relatively small, custom integrated proteogenomics database (iPtgxDB)(52)(53) for each of the 46,754 RefSeq assemblies.

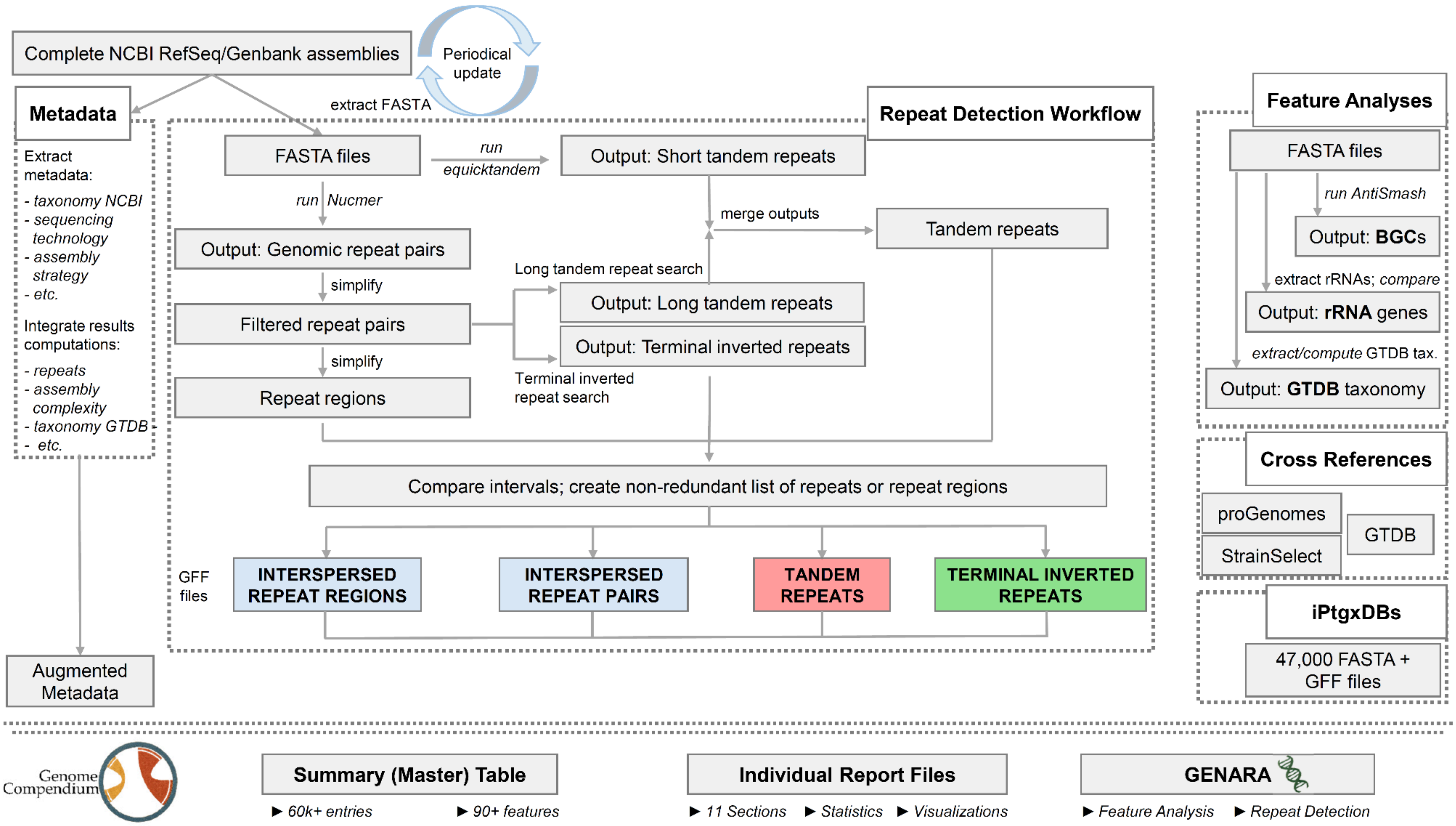
Complete NCBI RefSeq/Genbank assemblies
Periodical update
extract FASTA
Metadata
Extract metadata:
- taxonomy NCBI
- sequencing technology
- assembly strategy
- etc.
Integrate results computations:
- repeats
- assembly complexity
- taxonomy GTDB
- etc.
Augmented Metadata
Repeat Detection Workflow
FASTA files
run equicktandem
Output: Short tandem repeats
run Nucmer
Output: Genomic repeat pairs
merge outputs
Tandem repeats
simplify
Long tandem repeat search
Filtered repeat pairs
Output: Long tandem repeats
Output: Terminal inverted repeats
simplify
Terminal inverted repeat search
Repeat regions
Compare intervals; create non-redundant list of repeats or repeat regions
GFF files
INTERSPERSED REPEAT REGIONS
INTERSPERSED REPEAT PAIRS
TANDEM REPEATS
TERMINAL INVERTED REPEATS
Feature Analyses
FASTA files
run AntiSmash
Output: BGCs
extract rRNAs; compare
Output: rRNA genes
extract/compute GTDB tax.
Output: GTDB taxonomy
Cross References
proGenomes
GTDB
StrainSelect
iPtgxDBs
47,000 FASTA + GFF files
Genome Compendium
Summary (Master) Table
► 60k+ entries
► 90+ features
Individual Report Files
► 11 Sections
► Statistics
► Visualizations
GENARA
► Feature Analysis
► Repeat Detection

**Figure S3. Details of the repeat identification and classification approach.**

Overview of the six steps that are carried out to identify the different types of repeats, to group them into repeat regions and to thereby also simplify their visualization. The respective parameters used are shown in six boxes illustrated with gimmicks.

Using the FASTA sequences extracted for the complete genome assemblies from RefSeq and GenBank, Nucmer (1) and equicktandem are run to detect repeats and tandem repeats (length limit 10,000 bp). In Step1, all self-matches on the diagonal are removed. In Step2, the repeat pairs are simplified: fully contained repeats are swallowed, repeats with fuzzy ends are merged, as well as repeats that overlap by >95% of the length of the smaller repeat. The output file is a list of interspersed repeat pairs, which get further simplified to interspersed repeat regions in Step3, where fully contained repeats are swallowed and repeats overlapping to >95% of the smaller repeat are merged. In Step 4 and 5, long tandem repeats and terminal inverted and directed repeats are identified from the list of repeat pairs obtained in Step 2. The list of tandem repeat regions is combined with the equicktandem output. Interspersed repeat regions overlapping with a tandem and/or a terminal inverted repeat region for more than 95% are merged and deleted in Step 6. Subsequently, tandem regions are compared to the interspersed regions not deleted in the previous step; if the tandem region overlaps by >95% with an interspersed region, it gets merged with it. This is done to simplify the results and to prioritize tandem and terminal inverted repeat regions. The final lists of interspersed, tandem, and terminal inverted regions are written in three separate GFF files.

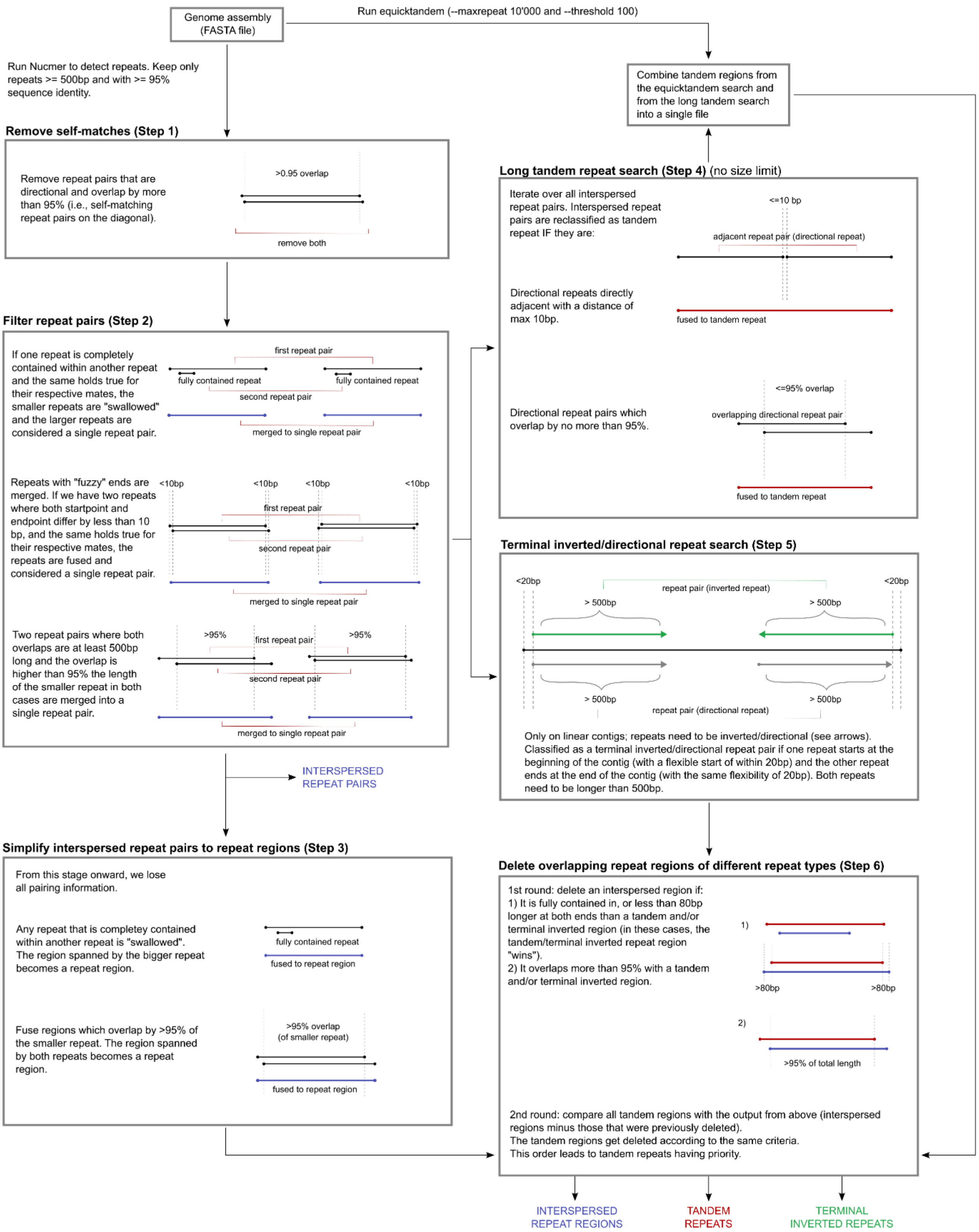
Genome assembly (FASTA file)
Run equicktandem (--maxrepeat 10'000 and --threshold 100)
Run Nucmer to detect repeats. Keep only repeats >= 500bp and with >= 95% sequence identity.
Combine tandem regions from the equicktandem search and from the long tandem search into a single file
Remove self-matches (Step 1)
Remove repeat pairs that are directional and overlap by more than 95% (i.e., self-matching repeat pairs on the diagonal).
>0.95 overlap
remove both
Long tandem repeat search (Step 4) (no size limit)
Iterate over all interspersed repeat pairs. Interspersed repeat pairs are reclassified as tandem repeat IF they are:
<=10 bp
adjacent repeat pair (directional repeat)
Directional repeats directly adjacent with a distance of max 10bp.
fused to tandem repeat
Directional repeat pairs which overlap by no more than 95%.
<=95% overlap
overlapping directional repeat pair
fused to tandem repeat
Filter repeat pairs (Step 2)
If one repeat is completely contained within another repeat and the same holds true for their respective mates, the smaller repeats are "swallowed" and the larger repeats are considered a single repeat pair.
first repeat pair
fully contained repeat
fully contained repeat
second repeat pair
merged to single repeat pair
Repeats with "fuzzy" ends are merged. If we have two repeats where both startpoint and endpoint differ by less than 10 bp, and the same holds true for their respective mates, the repeats are fused and considered a single repeat pair.
<10bp
<10bp
<10bp
<10bp
first repeat pair
second repeat pair
merged to single repeat pair
Two repeat pairs where both overlaps are at least 500bp long and the overlap is higher than 95% the length of the smaller repeat in both cases are merged into a single repeat pair.
>95%
first repeat pair
>95%
second repeat pair
merged to single repeat pair
Terminal inverted/directional repeat search (Step 5)
<20bp
repeat pair (inverted repeat)
<20bp
> 500bp
> 500bp
> 500bp
> 500bp
repeat pair (directional repeat)
Only on linear contigs; repeats need to be inverted/directional (see arrows). Classified as a terminal inverted/directional repeat pair if one repeat starts at the beginning of the contig (with a flexible start of within 20bp) and the other repeat ends at the end of the contig (with the same flexibility of 20bp). Both repeats need to be longer than 500bp.
INTERSPERSED REPEAT PAIRS
Simplify interspersed repeat pairs to repeat regions (Step 3)
From this stage onward, we lose all pairing information.
Any repeat that is completey contained within another repeat is "swallowed". The region spanned by the bigger repeat becomes a repeat region.
fully contained repeat
fused to repeat region
Fuse regions which overlap by >95% of the smaller repeat. The region spanned by both repeats becomes a repeat region.
>95% overlap (of smaller repeat)
fused to repeat region
Delete overlapping repeat regions of different repeat types (Step 6)
1st round: delete an interspersed region if:
1) It is fully contained in, or less than 80bp longer at both ends than a tandem and/or terminal inverted region (in these cases, the tandem/terminal inverted repeat region "wins").
2) It overlaps more than 95% with a tandem and/or terminal inverted region.
1)
>80bp
>80bp
2)
>95% of total length
2nd round: compare all tandem regions with the output from above (interspersed regions minus those that were previously deleted).
The tandem regions get deleted according to the same criteria.
This order leads to tandem repeats having priority.
INTERSPERSED REPEAT REGIONS
TANDEM REPEATS
TERMINAL INVERTED REPEATS

**Table S2. Enrichment analyses for various features.**

Enrichment analyses (hypergeometric tests) were carried out (mainly at the genus level, and for a few other taxonomic ranks) to explore whether assemblies from particular ranks were enriched in specific genome features such as repeats, 16S rRNA genes or biosynthetic gene clusters (see below). A multiple testing correction was carried out with the method of Holm, also called Holm-Bonferroni method (54). The results of these enrichment analyses are summarized on different sheets. **See separate Excel file**.

Individual sheets are described in detail in the legend sheet. They include **NoRepeats** - Assemblies without any repeats (441); **TIR** - Assemblies with TIRs (741); **One 16SrRNA** - Assemblies with exactly 1 16S rRNA copy (3,745); **<9 16SrRNA** - Assemblies with 1-8 16S rRNA copies (42,575); **>=9 16SrRNA** - Assemblies with 9 or more 16S rRNA copies (4,153); **>1 16S_identical** - Assemblies with multiple 16S rRNA copies that all have identical sequences per genome (14,742); **>1 16S_different** - Assemblies with multiple 16S rRNA copies where at least one copy per genome differs in sequence from the rest (28,241); **16SrRNA_Ident<99percent** - Assemblies with multiple 16S rRNA copies where at least one copy per genome has < 99% sequence identity (3,282); **16SrRNA_Ident<97percent** - Assemblies with multiple 16S rRNA copies where at least one copy per genome has < 97% sequence identity (meta-analysis, multiple taxonomic ranks); **Repeat>=30kb** - Assemblies with at least one repeat of 30kb or longer (2,724); **>=300Repeats** - Assemblies with over 300 repeats (648); **KorenClassII** - Assemblies for which at least 1 genome in a taxonomic rank family (3,228) is classified as class II. **>=10BGCs** - Assemblies of genomes with at least 10 BGCs (13,209). **Reference-based** - Assemblies for which at least one replicon was assembled with a reference-based strategy (1,351). **Repeats_GeneClasses** - First table - Enrichment in repeats across all analysed genomes using species as strata for features such as ribosomal protein genes (negative control), rRNA and transposase genes (positive controls) and then AMR genes overall plus 25 of 41 AMR subclasses that passed the inclusion threshold; Second table - Enrichment of AMR related genes in repeats within each genus using species as strata; only genera passing the thresholds are shown (86 of 654). Enrichment was assessed with specific filtering criteria, most importantly the presence of at least two features per genome. For details on how many assemblies were considered for the multiple testing correction for each subset, please see the respective sheet.

**Table S3. Prokaryotic genomes are more complex than reported previously.**

An analysis of the percentage of genomes in three assembly complexity classes from different analysis “snapshots” underlines that - as hypothesized earlier(55) - repeat-rich, complex to assemble class III genomes are significantly more prevalent than reported in 2018. Advances in sequencing technology and longer read lengths can explain this (see **Figure 3A**).

| **Datasets:** | **NAR* (~Feb2018)** | | **RefSeq (June2022)** | | **RefSeq (Dec2024)** | | **since NAR** | |
|---|---|---|---|---|---|---|---|---|
| Koren class | number | percent | number | percent | number | percent | number | percent |
| Class I | 4,842 | 54.9 | 14,233 | 51.0 | 23,410 | 50.1 | 18,576 | 49.0 |
| Class II | 781 | 8.9 | 2,109 | 7.6 | 3,228 | 6.9 | 2,447 | 6.4 |
| **Class III** | **3,195** | **36.2** | **11,565** | **41.4** | **20,116** | **43.0** | **16,927** | **44.6** |

*Approximation of a RefSeq dataset comparable to the time stamp Feb2018 from NCBI (in the original paper, we had analyzed GenBank assemblies, but since focus on the higher quality RefSeq genomes).

**Table S4. Association rule mining among BGCs.**

We performed an association rule mining among co-occurring genomic features, here BGCs, in order to identify combinations of BGCs that might also be predictive and characteristic for different bacterial genera. A content sheet describes which information is provided in the two additional sheets, an overview of enrichment tests (odds ratio >1, adjusted p-value < 0.01) and the uncovered strong association rules (close to 7000 with support>0.01, confidence>0.8 and lift>2). Selected relevant associations are described in the Results and Supplementary Results sections. **See separate Excel file**.

**Figure S4. Repeat region content of prokaryotic genomes.**

Visualization of the overall repeat region content (here in percent) of RefSeq and GenBank assemblies. GenBank contains a few genomes that represent duplications of the entire assembly (a repeat region value of 1), which are excluded from RefSeq.

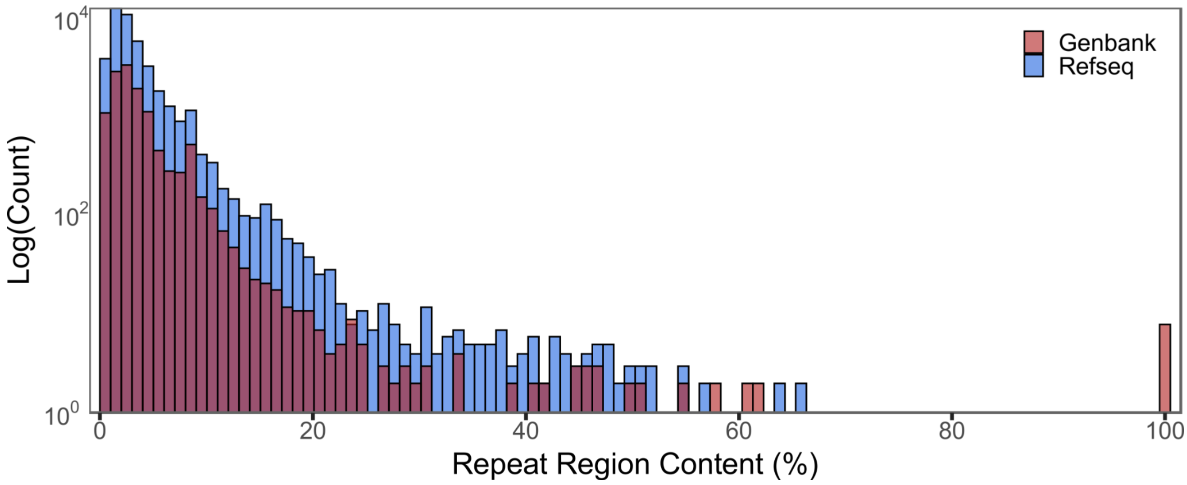

**Figure S5. Length distribution of different repeat types in subsets of RefSeq genomes.**

Across the plots below, distributions are shown for genome assemblies with (21,370) and without plasmids (25,384), alongside a central small, red boxplot representing the overall data without grouping. Outliers are indicated in white within the boxplot. **A.** Violin plots highlighting the repeat distributions for the entire respective subset dataset. **B.** A violin plot shows the length distribution of the respective longest copy of interspersed or tandem repeats found in 41,853 genomes that contain both types of repeats (89.5% of RefSeq assemblies). **C.** Violin plot showing the length distribution of different repeats (again, the longest repeat for a given subtype is used), here for the subset of 741 genomes that harbor terminal inverted repeats either on the chromosome and/or plasmid.

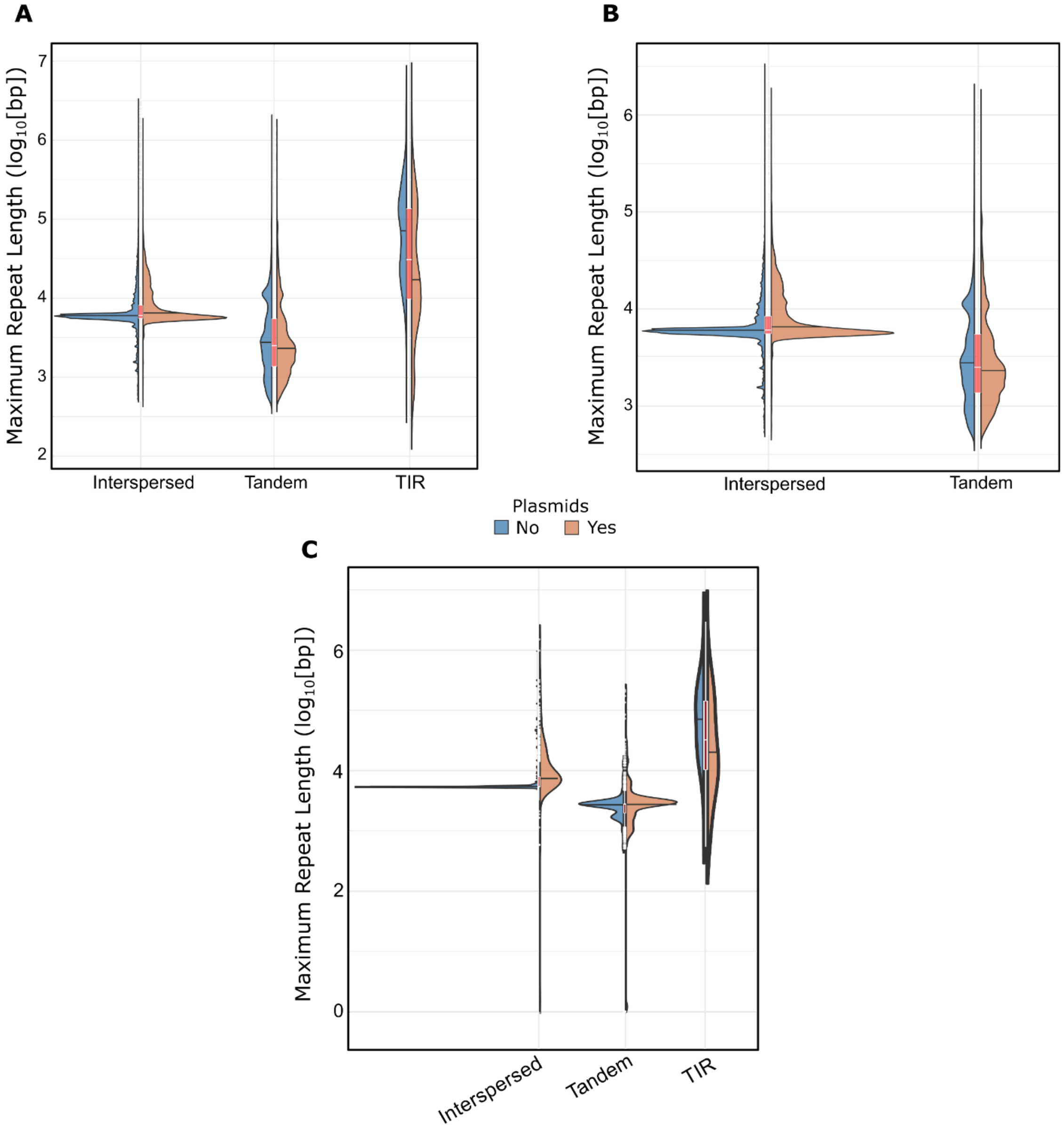

**Figure S6. Fraction of AMR genes found in repeats among ESKAPE pathogens.**

Analysis of the proportion of AMR genes identified in repeats in different species of six selected genera, here focusing on ESKAPE pathogens. The plot shows those species that meet the cut-off criteria for our analysis (see Methods) and the respective fraction of AMR genes identified by AMRFinderPlus that are associated with repeats. The specific ESKAPE species (ochre dots) all display a significant enrichment and include many genomes (dot size). Some species with exceptionally high proportions of AMR genes in repeats are identified (see additional labels), which typically only comprise few genomes, but may be worth-while to watch in the future.

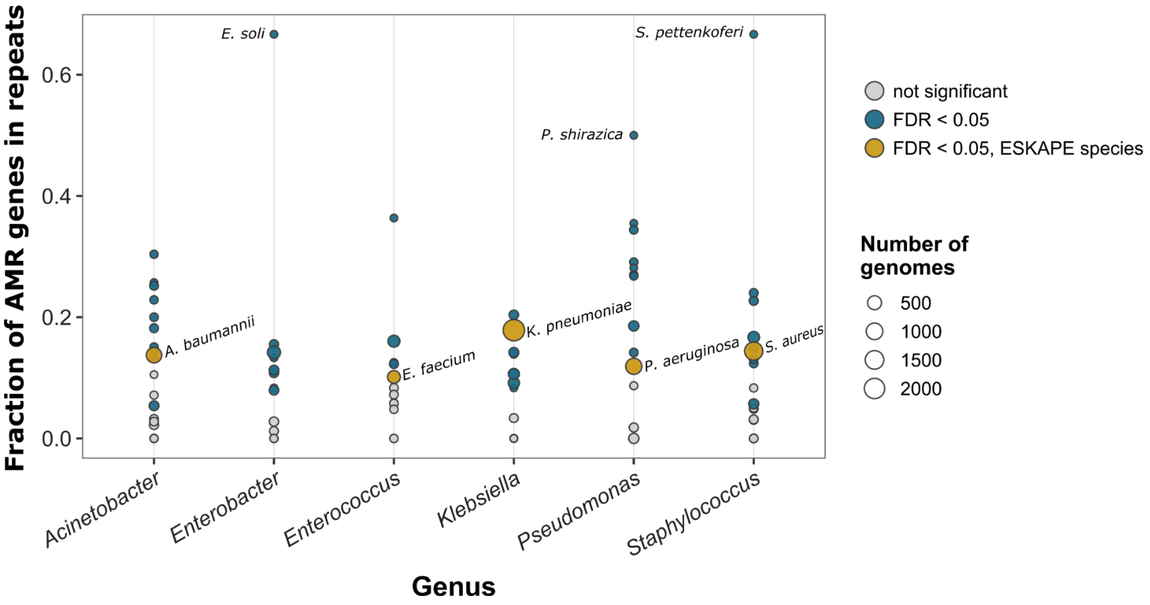

**Figure S7**. **Increasing percentage of difficult to assemble class III genomes over time**.

The cumulative percentage of complete genome sequences classified as class III (dark grey), i.e., the most difficult to assemble genomes with repeats well over 7 kbp in length, has been increasing over the years. By the end of December 2024, it amounted to 43.0%, indicating that earlier analyses have significantly underestimated the overall frequency of complex prokaryotic genomes. When considering genomes submitted up to the end of 2013, i.e., a time when the first genomes resolved by long read sequencing data started to appear (see **Figure 1D**), this percentage was only 27%.

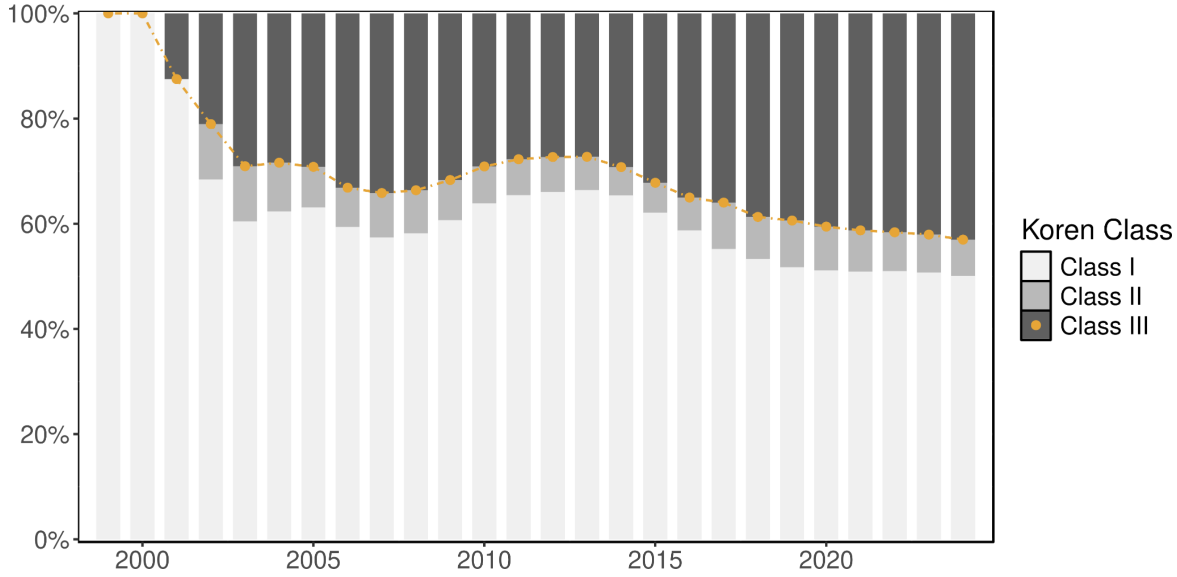

**Figure S8. Differences among taxonomy classifications from NCBI and GTDB.**

Overview of the differences in assignments for different taxonomic ranks between NCBI and GTDB. The percentage of entries for a given rank, for which the GTDB classification differs from that of the NCBI is shown in blue. Considering all ranks, overall, there is at least one different taxonomy assignment for 64% of the RefSeq assemblies.
Missing in GTDB: when the name of a taxon is present in NCBI but missing in GTDB. Missing in NCBI: when the name of a taxon is missing in NCBI but present in GTDB.
Missing in both: a taxon name is absent in both NCBI and GTDB. The respective counts of entries (assemblies) missing only in GTDB, only in NCBI RefSeq or missing in both are also summarized in tabular format below.

| | Domain | Phylum | Class | Order | Family | Genus | Species |
|---|---|---|---|---|---|---|---|
| **Missing in GTDB** | 0 | 0 | 0 | 1 | 1 | 6 | 102 |
| **Missing in NCBI** | 0 | 9 | 514 | 76 | 187 | 26 | 5,410 |
| **Missing in both** | 0 | 0 | 0 | 0 | 0 | 0 | 206 |

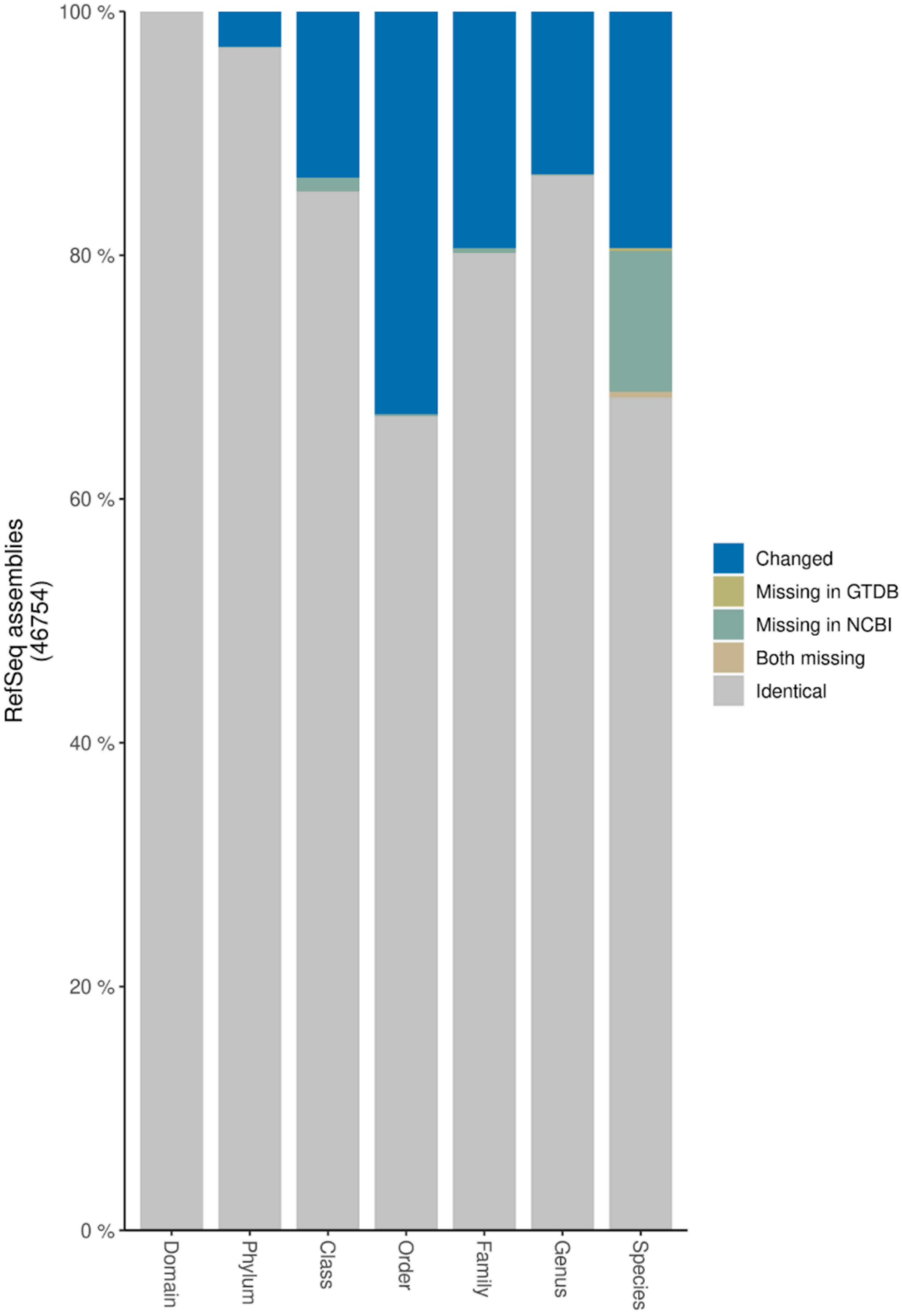

100 %
80 %
60 %
40 %
20 %
0 %
RefSeq assemblies
(46754)
Domain
Phylum
Class
Order
Family
Genus
Species
Changed
Missing in GTDB
Missing in NCBI
Both missing
Identical

**Figure S9. Exploring the increasing taxonomic coverage of RefSeq genomes over time.**

**A**. Number of distinct novel taxonomic ranks added per year. Data was fitted with a generalized linear model directly when plotting the data with the R package geom_smooth. For the higher taxonomic ranks phylum, class and order, the linear fit shows a decreasing yearly increase of novel entries at the respective taxonomic rank. This trend is reversed for the taxonomic ranks family, genus and species, where a yearly increase can be noted (see **Figure 5A**). **B**. A metacoder plot for the order Hyphomicrobiales (see also **Figure 5 C, D**) visualizes which species have been newly added since the NAR dataset (in ochre). Amongst others, this order includes important plant growth promoting bacteria (PGPB). The plot implies the genera *Rhizobium* and *Bradyrhizobium* as hotspots. In contrast, only two new species were added for example for the genus *Brucella* (*Brucella intermedia* and *Brucella pseudointermedia*, highlighted in red).

**A.**

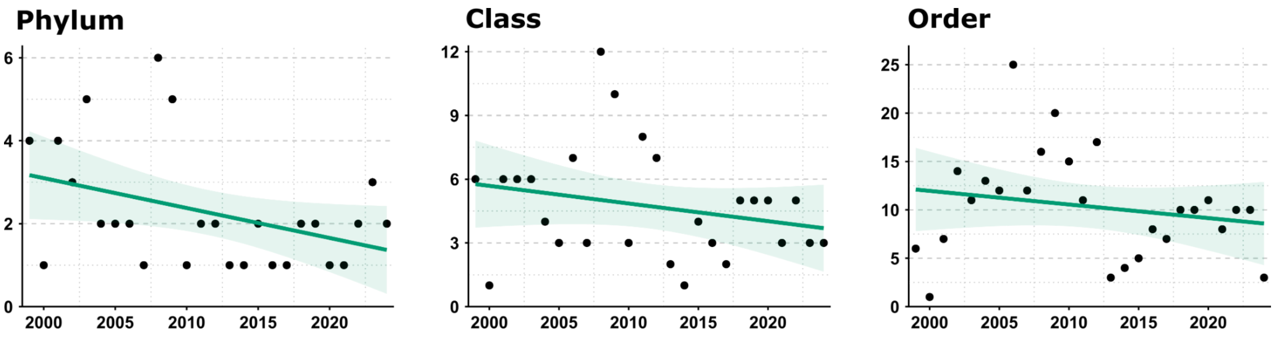

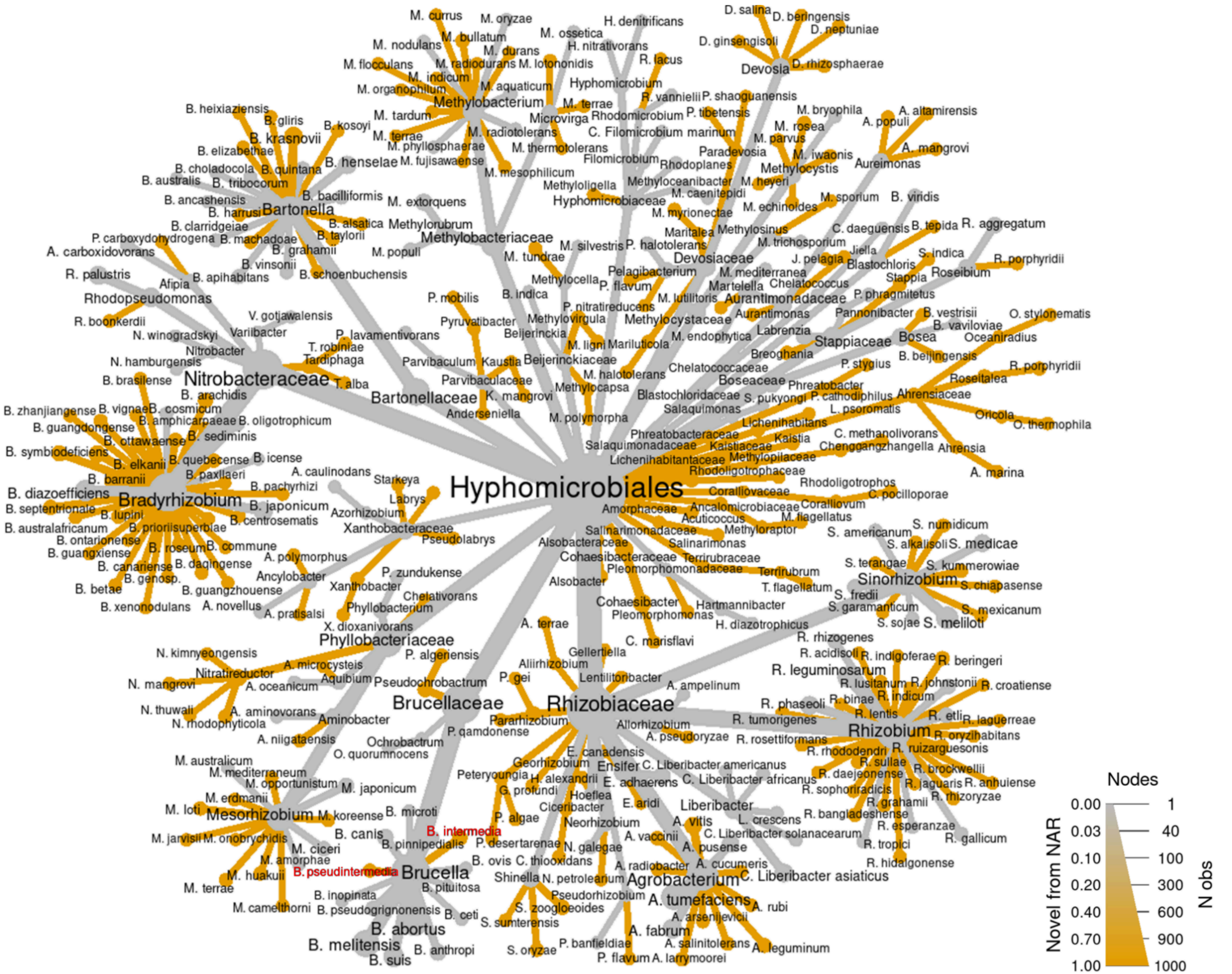
Hyphomicrobiales
Rhizobiaceae
Brucellaceae
Phyllobacteriaceae
Xanthobacteraceae
Nitrobacteraceae
Bartonellaceae
Methylobacteriaceae
Hyphomicrobiaceae
Devosiaceae
Aurantimonadaceae
Methylocystaceae
Stappiaceae
Boseaceae
Ahrensiaceae
Beijerinckiaceae
Parvibaculaceae
Blastochloridaceae
Salaquimonadaceae
Phreatobacteraceae
Lichenihabitantaceae
Kaistiaceae
Methylopilaceae
Rhodoligotrophaceae
Cohaesibacteraceae
Pleomorphomonadaceae
Alsobacteraceae
Amorphaceae
Salinarimonadaceae
Terrirubraceae
Corallivaceae
Ancalomicrobiaceae
Bradyrhizobium
Bartonella
Methylobacterium
Devosia
Sinorhizobium
Rhizobium
Agrobacterium
Brucella
Mesorhizobium
Liberibacter
Ensifer
B. intermedia
B. pseudintermedia
B. abortus
B. melitensis
B. suis
B. canis
B. ovis
B. ceti
B. pinnipedialis
B. microti
B. inopinata
B. pseudogrignonensis
B. anthropi
B. pituitosa
B. henselae
B. quintana
R. leguminosarum
A. tumefaciens
A. fabrum
S. meliloti
S. medicae
C. Liberibacter asiaticus
C. Liberibacter americanus
C. Liberibacter africanus
C. Liberibacter solanacearum
Nodes
0.00
0.03
0.10
0.20
0.40
0.70
1.00
1
40
100
300
600
900
1000
Novel from NAR
N obs

B.

**Figure S10. Top 20 species in terms of number of assemblies.**

The list of 20 species with the highest number of assemblies is dominated by pathogens, including the medically relevant ESKAPE pathogens (*Enterococcus faecium, Staphylococcus aureus, Klebsiella pneumoniae, Acinetobacter baumannii, Pseudomonas aeruginosa, Enterobacter species)*. In the plot below, we show the overall count of the top 20 species, and further differentiate the respective added counts for the time frames up to 2010 (dark green), 2010-2014 (light green), 2015-2019 (cream) and 2020-2024 (dark gray bar).

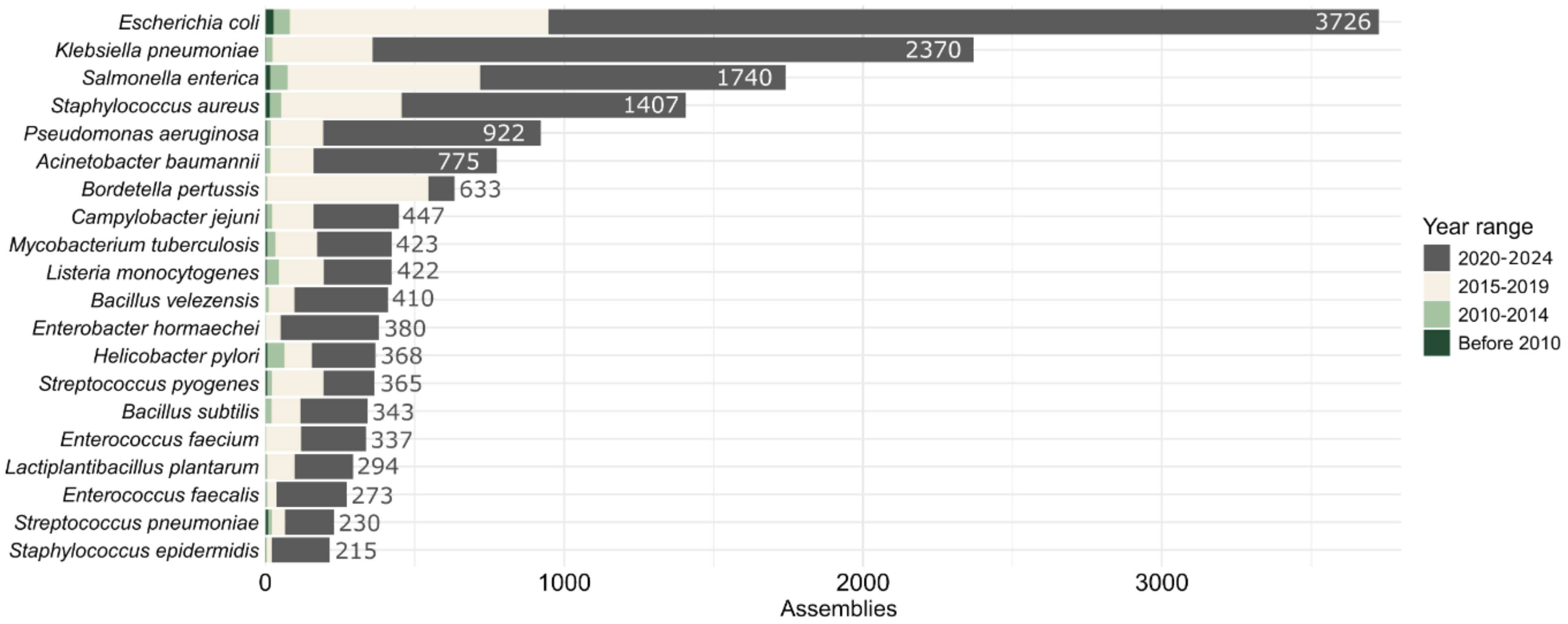

**Figure S11. Increased taxonomic coverage for the order Lactobacillales.**

**A.** Several genera are labeled on the ggtree plot for the order Lactobacillales, i.e., genera for which a larger number of distinct species was observed (see outer ring and the legend) or where many novel genera or higher taxonomic ranks (here the three families Lactobacillaceae, Carnobacteriaceae, Aerococcaceae) were implied by the novel species that have been added since the analysis in 2018. The inner circle shows the percentage of novelty compared to the NAR2018 time point, the outer circle the number of distinct species added for a given genus. Two families are labeled, i.e., Carnobacteriaceae and Aerococcaceae (bold text), for which new species from several distinct genera were completely sequenced. This is also true for the family Lactobacillaceae (see part of the plot from roughly 4 o'clock to almost 10 o'clock of Figure A, where 9 genera with novel species were added). **B.** Metacoder plot for the order Lactobacillales, which, amongst others, includes important bacteria used in fermentation of foods and cheeses. Again, species that have been newly added since the NAR dataset are shown in ochre. The plot illustrates that the genera *Streptococcus*, *Lactobacillus* are nodes where several new species were added and that the family Aerococcaceae has become much better described with many new branches added. In contrast, only two new species are added for the genus *Leuconostoc* (*L. falkenbergense, L. pseudomesenteroides*, highlighted in red).

**A.**

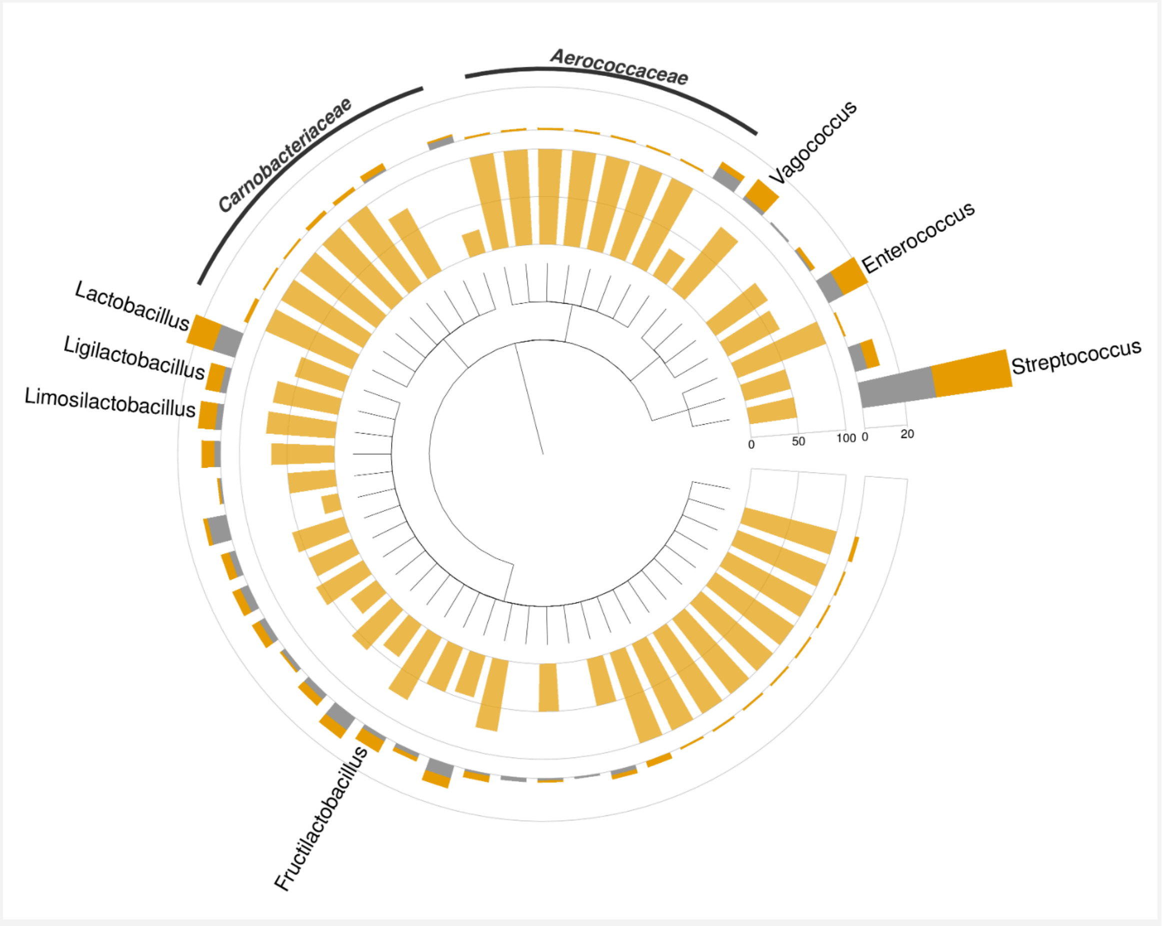

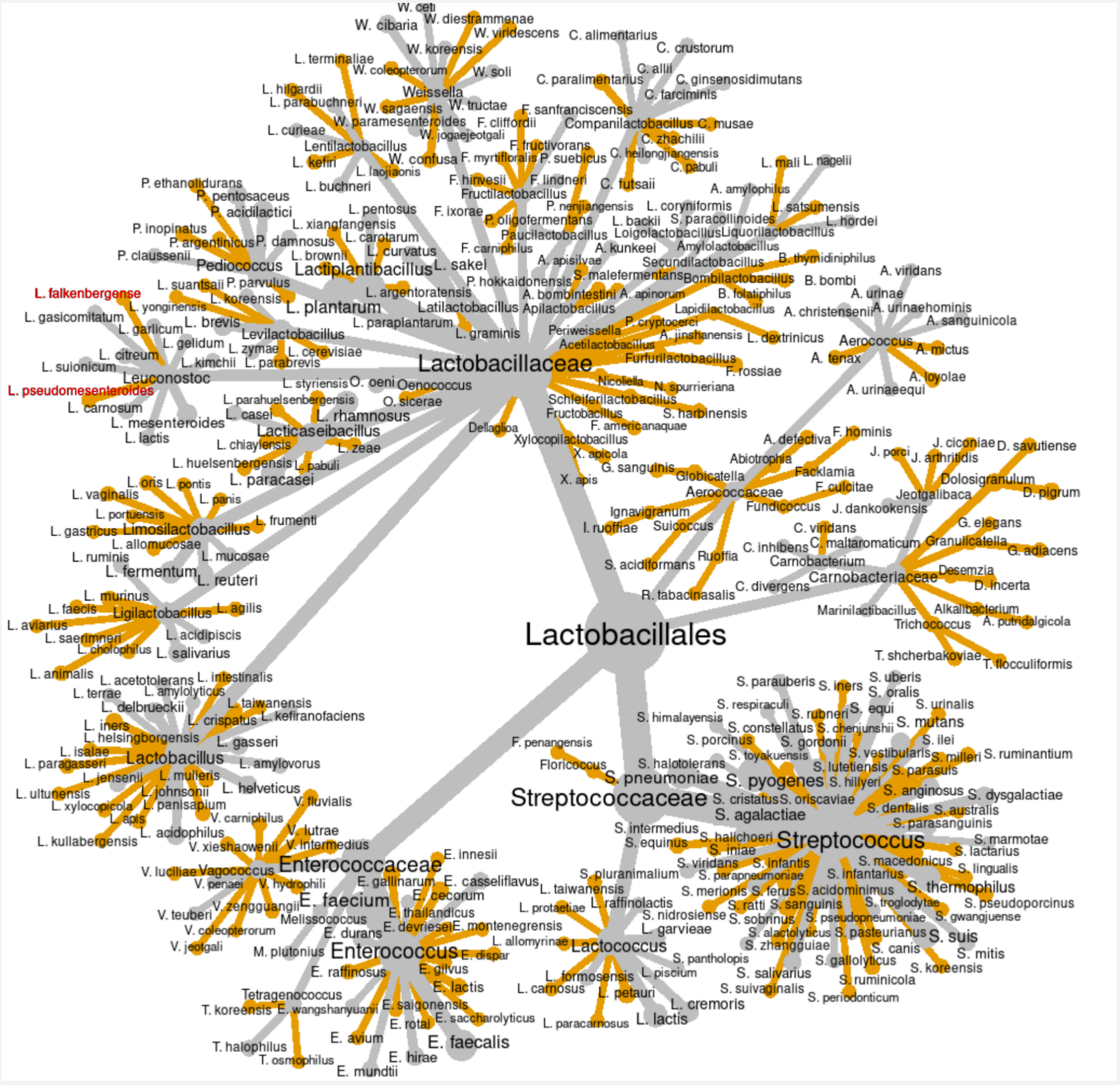
Lactobacillales
Lactobacillaceae
Streptococcaceae
Enterococcaceae
Aerococcaceae
Carnobacteriaceae
W. ceti
W. cibaria
W. diestrammenae
W. viridescens
W. koreensis
W. coleopterorum
W. soli
Weissella
W. sagaensis
W. tructae
W. paramesenteroides
W. jogaejeotgali
W. confusa
C. alimentarius
C. crustorum
C. allii
C. paralimentarius
C. ginsenosidimutans
C. farciminis
Companilactobacillus
C. musae
C. zhachilii
C. heilongjiangensis
C. pabuli
C. futsaii
F. sanfranciscensis
F. cliffordii
F. fructivorans
F. myrtifloralis
P. suebicus
F. hinvesii
F. lindneri
Fructilactobacillus
F. ixorae
P. nenjiangensis
P. oligofermentans
Paucilactobacillus
F. carniphilus
L. terminaliae
L. hilgardii
L. parabuchneri
L. curieae
Lentilactobacillus
L. kefiri
L. laojiaonis
L. buchneri
L. pentosus
L. xiangfangensis
L. carotarum
L. curvatus
L. brownii
Lactiplantibacillus
L. sakei
L. argentoratensis
L. plantarum
L. paraplantarum
Latilactobacillus
L. graminis
P. ethanolidurans
P. pentosaceus
P. acidilactici
P. inopinatus
P. argentinicus
P. damnosus
P. claussenii
Pediococcus
P. parvulus
L. suantsaii
L. falkenbergense
L. yonginensis
L. koreensis
L. gasicomitatum
L. garlicum
L. brevis
Levilactobacillus
L. gelidum
L. zymae
L. cerevisiae
L. citreum
L. suionicum
L. kimchii
L. parabrevis
Leuconostoc
L. pseudomesenteroides
L. carnosum
L. mesenteroides
L. lactis
L. styriensis
O. oeni
Oenococcus
L. parahuelsenbergensis
O. sicerae
L. casei
L. rhamnosus
Lacticaseibacillus
L. chiayiensis
L. zeae
L. huelsenbergensis
L. pabuli
L. paracasei
Dellaglioa
L. oris
L. pontis
L. vaginalis
L. panis
L. portuensis
L. frumenti
L. gastricus
Limosilactobacillus
L. allomucosae
L. ruminis
L. mucosae
L. fermentum
L. reuteri
L. murinus
L. faecis
Ligilactobacillus
L. agilis
L. aviarius
L. saerimneri
L. acidipiscis
L. cholophilus
L. salivarius
L. animalis
L. acetotolerans
L. intestinalis
L. terrae
L. amylolyticus
L. delbrueckii
L. taiwanensis
L. iners
L. crispatus
L. kefiranofaciens
L. helsingborgensis
L. isalae
L. gasseri
L. paragasseri
Lactobacillus
L. amylovorus
L. jensenii
L. mulleris
L. ultunensis
L. johnsonii
L. helveticus
L. xylocopicola
L. panisapium
L. apis
L. acidophilus
L. kullabergensis
L. corynifornis
L. backii
S. paracollinoides
Loigolactobacillus
Liquorilactobacillus
A. kunkeei
Amylolactobacillus
A. apisilvae
Secundilactobacillus
P. hokkaidonensis
S. malefermentans
Bombilactobacillus
A. bombintestini
A. apinorum
Apilactobacillus
B. folatiphilus
Lapidilactobacillus
Periweissella
P. cryptocerci
A. jinshanensis
Acetilactobacillus
L. dextrinicus
Furfurilactobacillus
F. rossiae
Nicoliella
N. spurrieriana
Schleiferilactobacillus
Fructobacillus
S. harbinensis
F. americanaquae
Xylocopilactobacillus
X. apicola
G. sanguinis
X. apis
L. mali
L. nagelii
A. amylophilus
L. satsumensis
L. hordei
B. thymidiniphilus
B. bombi
A. viridans
A. urinae
A. christensenii
A. urinaehominis
A. sanguinicola
Aerococcus
A. tenax
A. mictus
A. loyolae
A. urinaeequi
A. defectiva
F. hominis
Abiotrophia
Facklamia
Globicatella
F. culicitae
Fundicoccus
Ignavigranum
I. ruoffiae
Suicoccus
S. acidiformans
Ruoffia
R. tabacinasalis
J. porci
J. ciconiae
J. arthritidis
D. savutiense
Dolosigranulum
D. pigrum
Jeotgalibaca
J. dankookensis
C. viridans
C. inhibens
C. maltaromaticum
Carnobacterium
C. divergens
G. elegans
Granulicatella
G. adiacens
Desemzia
D. incerta
Marinilactibacillus
Alkalibacterium
A. putridalgicola
Trichococcus
T. shcherbakoviae
T. flocculiformis
S. parauberis
S. iners
S. uberis
S. oralis
S. respiraculi
S. himalayensis
S. rubneri
S. equi
S. urinalis
S. constellatus
S. chenjunshii
S. mutans
S. porcinus
S. gordonii
S. ilei
S. toyakuensis
S. vestibularis
S. milleri
S. ruminantium
F. penangensis
S. halotolerans
S. lutetiensis
S. parasuis
Floricoccus
S. pneumoniae
S. pyogenes
S. hillyeri
S. anginosus
S. dysgalactiae
S. cristatus
S. oriscaviae
S. dentalis
S. australis
S. agalactiae
S. parasanguinis
S. intermedius
S. halichoeri
Streptococcus
S. marmotae
S. equinus
S. iniae
S. lactarius
S. viridans
S. infantis
S. macedonicus
S. parapneumoniae
S. infantarius
S. lingualis
S. pluranimalium
S. acidominimus
S. thermophilus
S. merionis
S. ferus
S. ratti
S. sanguinis
S. troglodytae
S. pseudoporcinus
S. nidrosiense
S. sobrinus
S. pseudopneumoniae
S. gwangjuense
S. alactolyticus
S. pasteurianus
S. suis
S. zhangguiae
S. canis
S. mitis
S. pantholopis
S. gallolyticus
S. koreensis
S. salivarius
S. ruminicola
S. suivaginalis
S. periodonticum
L. taiwanensis
L. protaetiae
L. raffinolactis
L. garvieae
L. allomyrinae
Lactococcus
L. formosensis
L. piscium
L. carnosus
L. petauri
L. cremoris
L. paracarnosus
L. lactis
V. fluvialis
V. carniphilus
V. lutrae
V. xieshaowenii
V. intermedius
Enterococcaceae
E. innesii
V. luciliae
Vagococcus
V. penaei
V. hydrophili
E. gallinarum
E. casseliflavus
V. zengguangii
E. faecium
E. cecorum
V. teuberi
Melissococcus
E. thailandicus
V. coleopterorum
E. durans
E. devriesei
E. montenegrensis
V. jeotgali
M. plutonius
Enterococcus
E. dispar
E. raffinosus
E. gilvus
E. lactis
Tetragenococcus
T. koreensis
E. wangshanyuanii
E. saigonensis
E. saccharolyticus
E. rotai
E. faecalis
T. halophilus
E. avium
T. osmophilus
E. hirae
E. mundtii
B

**Figure S12. Frequency distribution of protocluster classes across all RefSeq assemblies with valid genus-level assignments.**

Genome sequences were tokenized into binary presence-absence features representing distinct BGC product types. The distribution highlights the predominance of nonribosomal peptide (nrps) and ribosomally synthesized and post-translationally modified peptide (RiPP) clusters, as well as frequently observed terpene-precursor protoclusters. These high-frequency BGC classes form the dominant baseline from which subsequent co-occurrence rules were derived.

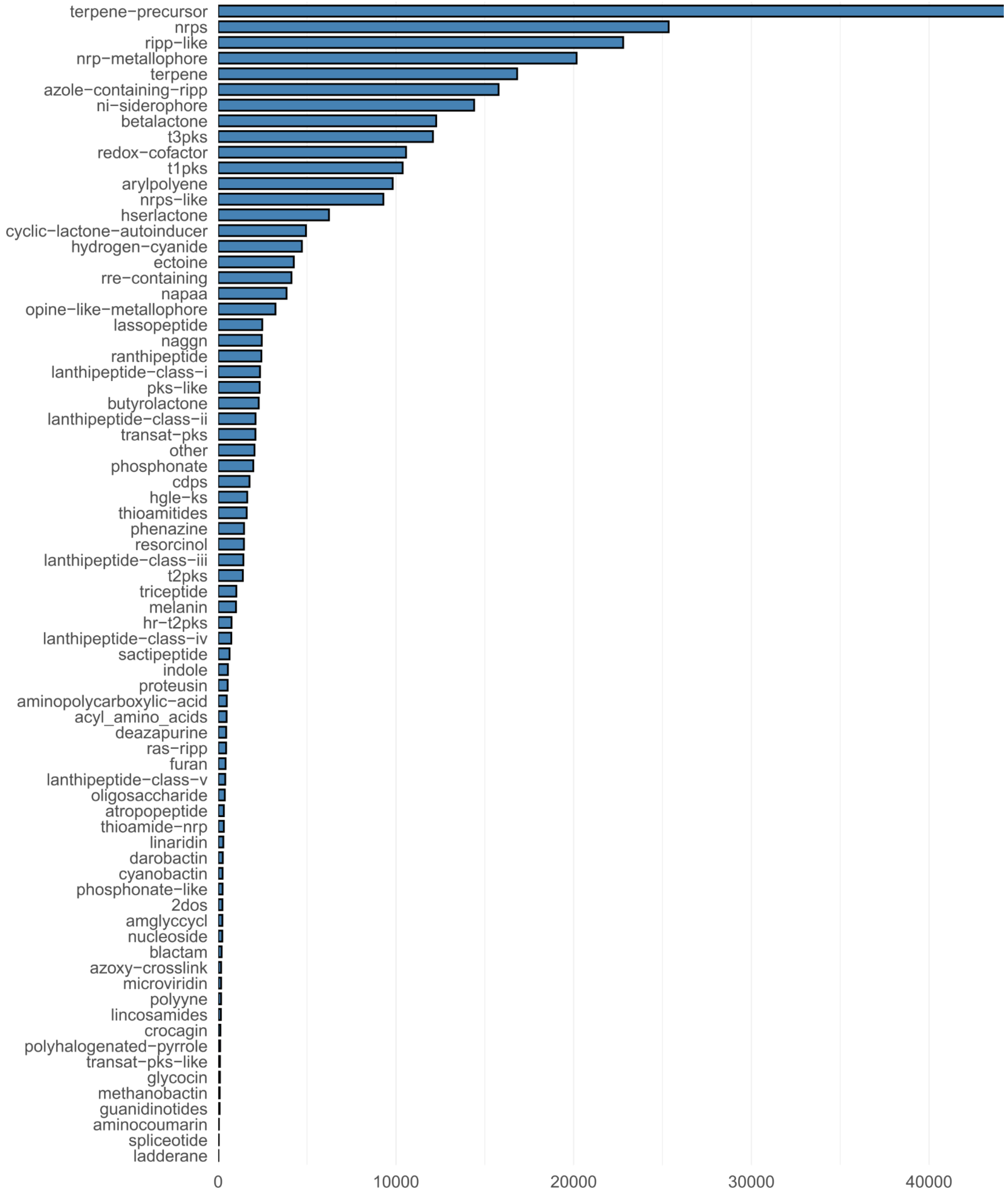

**Figure S13**. **Co-occurrence rule networks highlight genus-specific relations.**

Robust co-occurrence rules among protocluster types identified via association rule mining. Rules with high lift (>10), confidence (>0.95), and support (>1%) reveal strong dependencies between specific protoclusters (only 54 rules between 16 BGCs), such as opine-like metallophore, hydrogen cyanide, phenazine, and naggn clusters that were found to have characteristic associations for the genus *Pseudomonas (see* **Supplementary Table S4)**. The size of each rule reflects its support value. Rules are colored based on the colors assigned to their input BGCs. Links from rules to BGCs are emphasized (black) to distinguish them from links connecting BGCs to rules (light gray).

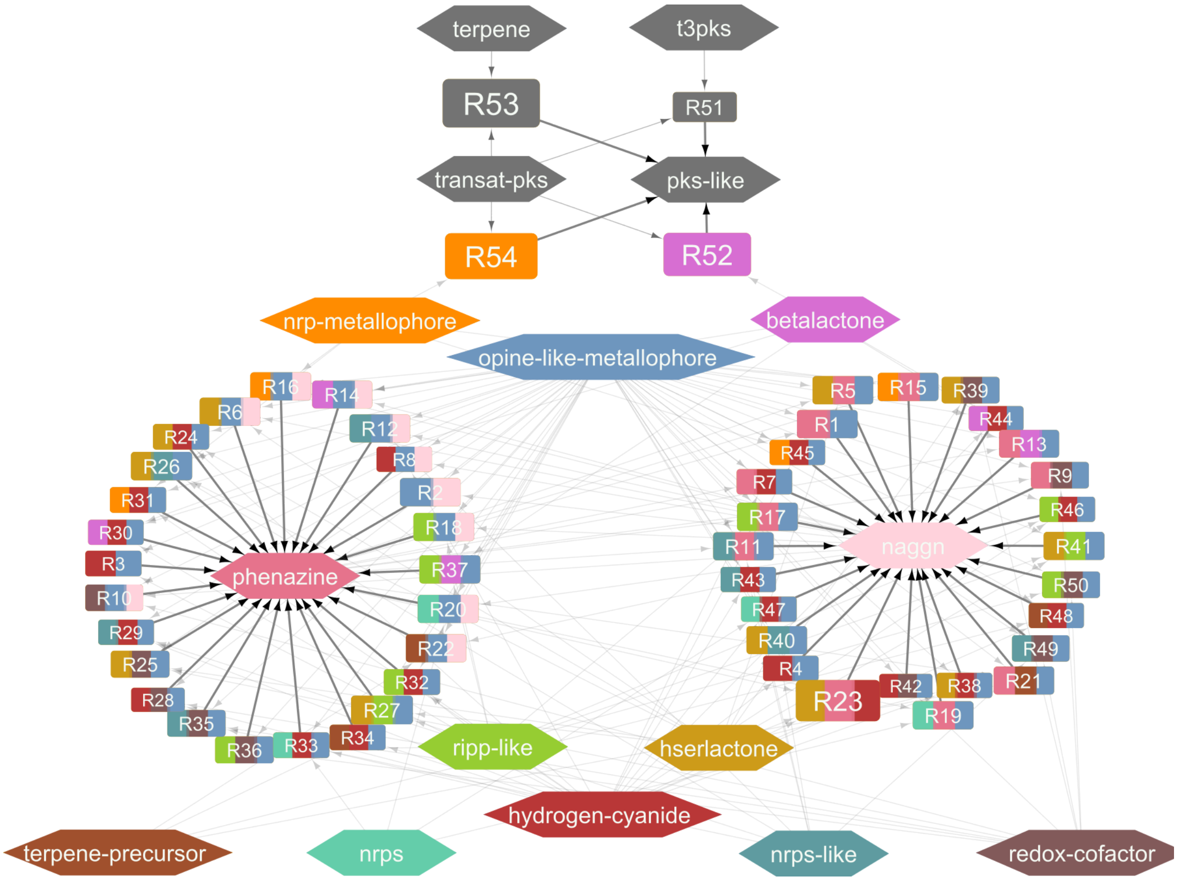

**Figure S14. Increase of distinct strains procurable from Bioresources listed by StrainSelect.**

The Bioresource centers (abbreviations from StrainSelect(51)), here ordered with respect to the number of available strains (from highest to lowest), starting with the German collection of microorganisms at the Leibniz Center (DSM), followed by the American Type Culture Collection (ATCC) and so on. **A**. In total, the BRCs provide access to 3,935 distinct stains. **B**. This corresponded to a total of 3,073 distinct species.

For these plots, only bioresource centers that added at least 10 unique additional strains to the collection were considered.

**A.**

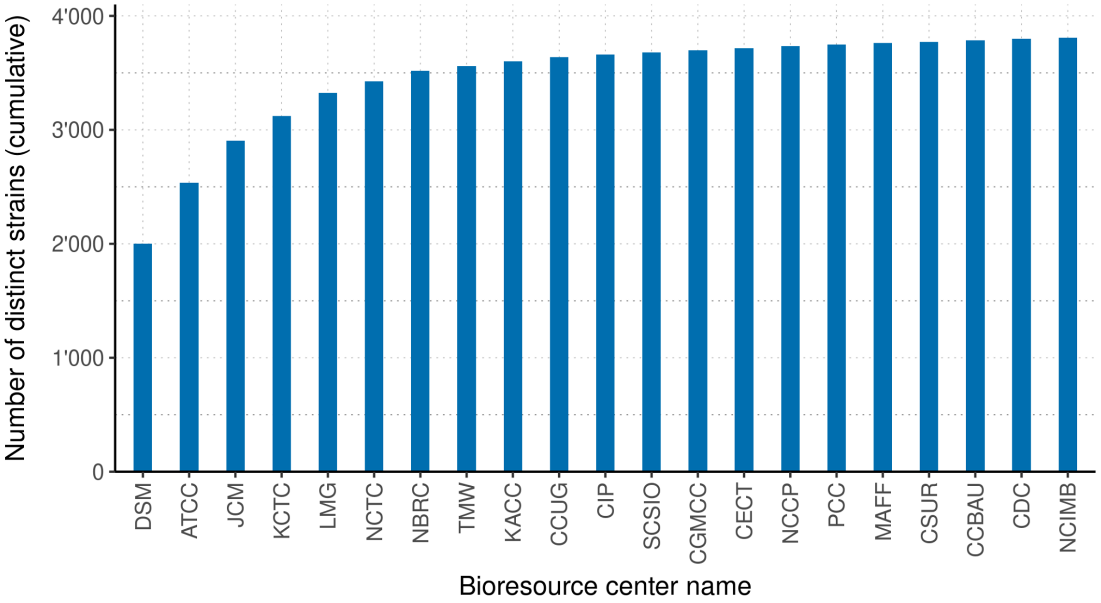

**B.**

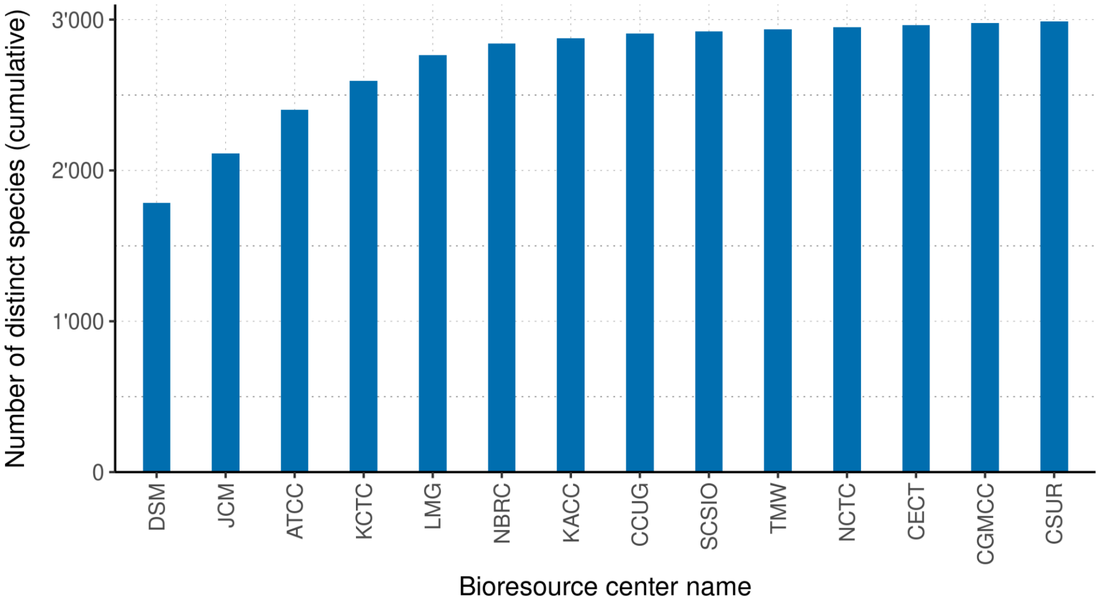

**Figure S15. Schematic overview and relevant aspects for the creation of iPtgxDBs.**

**A**. Integrated proteogenomics databases integrate different annotation resources for an identical genome sequence. These can include those from reference genome annotation centers (e.g. NCBI RefSeq, blue), ab initio predictions (e.g. from Prodigal, green), and other prediction sources (grey). Searching proteomics data against iPtgxDBs allows to identify experimental evidence for expressed pseudogenes, different start sites than annotated and novel, so far not annotated CDS (see also https://iptgxdb.expasy.org/ for more details about the concepts presented below). **B**. The different annotation resources are integrated in a hierarchical manner, starting with the most credible annotation resource as anchor annotations (here from NCBI RefSeq), and then integrating other annotations in a stepwise manner. **C**. Standard iPtgxDBs are very large (often more than 100,000 entries) as they integrate the in silico ORF predictions of a more advanced version of a six-frame translation (allowing alternative start codons and a user-defined length threshold of novel CDS) to capture almost the entire protein coding potential of a prokaryotic genome (52). Small, custom iPtgxDBs that replace the large number of in silico ORFs with smaller sets of potential candidates, here from the GMSC, have benefits for the proteomics searches (53, 56), as they only contain a median of 4,400 entries. **D**. Box plots of the number of annotated CDS for each of the included annotation sources, showing that RefSeq and Prodigal predict a median of roughly 4,000 CDS per genome while GMSC predicts ~400 SEPs. **E**. Hierarchical reduction of redundancy during the iPtgxDB creation reveals that Prodigal predicts a median of 347 novel proteins per genome compared to the RefSeq annotation, while GMSC additionally predicts a median of 25 novel SEPs that are neither found in the RefSeq nor the Prodigal annotations.

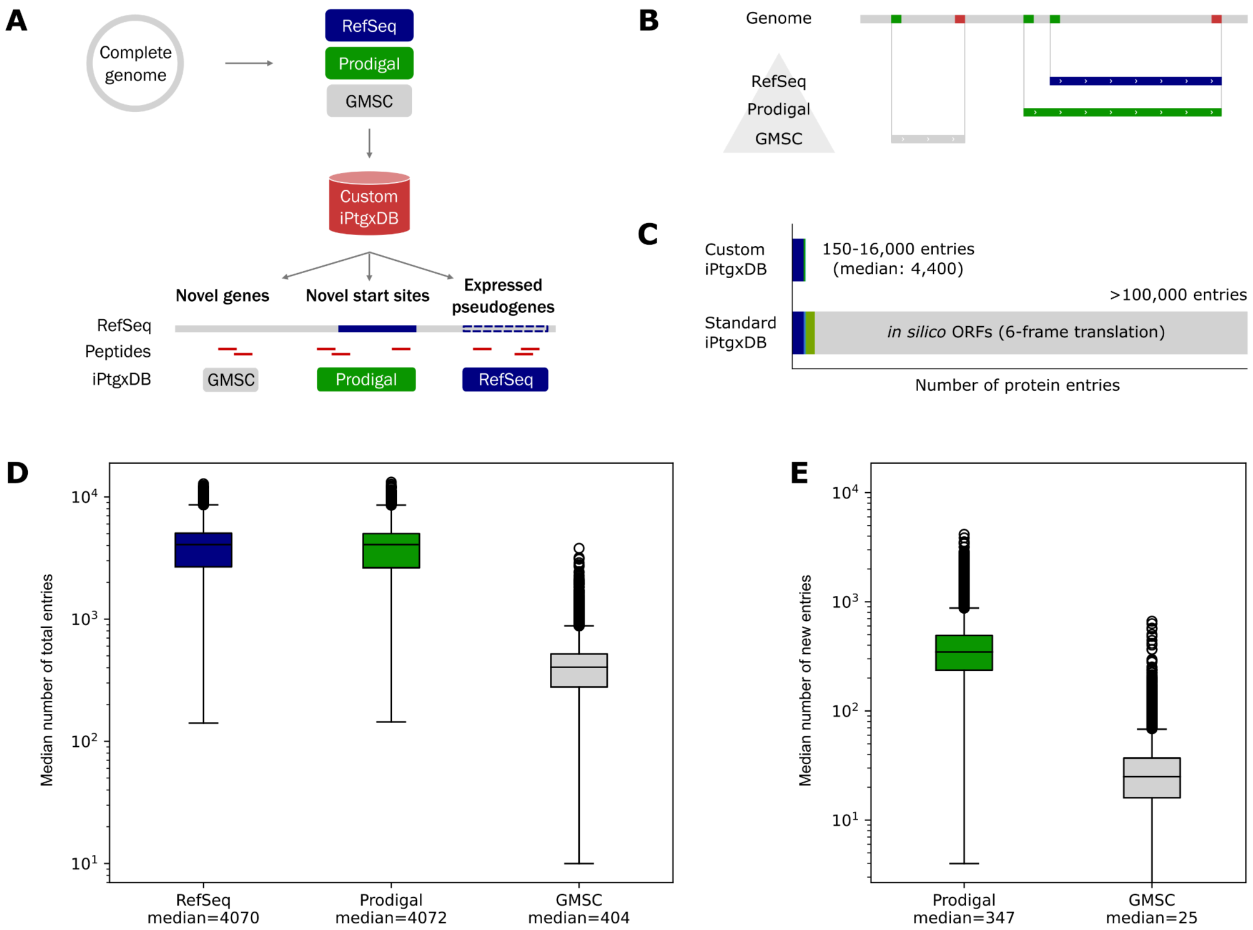
